# Early Emergence of Projection-subtype fate-restricted Radial Glial Progenitors Orchestrates Neocortical Neurogenesis

**DOI:** 10.1101/2025.05.07.652665

**Authors:** I. Varela-Martínez, A. Villalba, J. García-Marqués, S. Hippenmeyer, M. Nieto

## Abstract

Radial glial progenitors (RGPs) generate all projection neurons (PNs) in the cerebral cortex through incompletely understood processes. Herein, we combine Mosaic Analysis with Double Markers (MADM)-based clonal analysis at embryonic days 12.5 and 13.5 with early postnatal callosal tracing to reveal a lineage progression that challenges the inside-outside model of cortical development and the conventional view of an invariable sequence of asymmetric neurogenic divisions. Our data demonstrate that early multipotent RGPs generate all extra-telencephalic (ET) and intra-telencephalic (IT) PNs across all layers through parallel sublineages and the random specification, during the earliest neurogenic divisions, of fate-restricted daughter RGPs. While the neuronal production of the parental multipotent RGPs consists of small ET-PN or IT-PN outputs, fate-restricted RGPs produce larger translaminar outputs spanning deep and upper layers of only IT-PNs, the predominant mammalian PN subtype. We further show that the emergence of IT-PN fate-restricted RGPs also leads to quantitatively and temporally stereotyped neurogenesis population-wise.

## INTRODUCTION

The neocortex, a hallmark of mammalian evolution, is crucial for executing higher-order functions. Central to these functions are complex cortical networks of excitatory projection neurons (PNs). During development, radial glial progenitors (RGPs) generate all cortical PNs through a multistep process (1–5). It is thought that the neocortex evolved from an ancestral pallium through variations in developmental mechanisms and telencephalic progenitor cells involved in RGP lineages. However, the underlying mechanisms driving mammalian neurogenesis from RGPs remain only partially understood.

Mosaic Analysis with Double Markers (MADM) in mice has provided a robust framework for understanding neurogenesis (6, 7). MADM-based clonal analysis of RGP lineages has revealed that RGPs produce all PNs of the cerebral cortex following an organized program of proliferation and differentiation (8–10). First, during an amplification or symmetric phase, RGPs undergo a defined number of symmetric proliferative divisions, expanding the progenitor pool. Around embryonic day (E)12, RGPs transition into a neurogenic state and divide asymmetrically to allegedly produce PN subtypes sequentially for the different layers (11, 12). Once neurogenesis is completed, one out of six neurogenic RGPs switches to gliogenesis, while the rest undergo terminal divisions or differentiate (8–10). Despite this well-established framework, lineage progression during the neurogenic phase is mostly unknown (13–16) (17–21) (8, 22, 23).

A critical aspect of understanding cortical neurogenesis lies in finding a robust method for projection neuron (PN) classification. The conventional criterion in lineage studies relies on layer position as a readout of PN diversity; however, layer position alone does not adequately define PN identity, as different PN subtypes occupy the same cortical layer. Another useful criterion to describe PN identity is axonal patterns, which divide PNs into two major categories: intra-telencephalic (IT) PNs, which connect to different cortical regions and/or the basal telencephalon and are located across all layers (24, 25), and extra-telencephalic (ET) PNs, which target subcortical areas outside the telencephalon such as the thalamus, pons, and the spinal cord, and are only found in the deep layers (DLs) (26–29). The difficulty in attempting a projection-based classification is that no straightforward tracing method can comprehensively capture the full axonal diversity of adult IT-PNs and ET-PNs. In contrast, developing PNs share exuberant axonal projections that can be leveraged to identify the projection subtypes of adult PNs (30–32). In particular, studies from our laboratory have demonstrated that the populations of layer 2/3 and layer 4 PNs (UL PNs) project all through the corpus callosum (CC) by default and at early developmental stages (33). Such display of interhemispheric projections coincides with the transient expression of a callosal projection neuron (CPN) identity along a dynamic process of postnatal postmitotic differentiation (34) and many L2/3 and L4 callosal projections are developmentally eliminated. Nevertheless, since UL PNs comprise pure IT-PN populations, the data indicates that developmental callosal axons are a signature of immature IT-PN identity. Notably, numerous PNs in the DLs also display developmental callosal axons but whether these projections identify IT-PN identities has not been examined (33, 35).

Herein, we perform an RGP lineage analysis leveraging the early projection patterns of developing PNs. We first demonstrate that injections of fluorescent axonal tracers in the CC performed at early postnatal stages serve to distinguish extra-telencephalic (ET) from intra-telencephalic (IT) PNs. Then, by combining early callosal tracing with classical birth-dating strategies, we demonstrate that ET-PN and IT-PN populations are born sequentially. Namely, many ET-PNs and IT-PNs sharing the same layers are born asynchronously. These data indicated that that projection identity should be a cardinal decision that hierarchically predates layer identity at the time of specification from RGP progenitors. Moreover, given the established knowledge, we expected that RGPs would display “ET-PN” and “IT-PN” competence sequentially along their lineage progression. In contrast, Mosaic Analysis with Double Markers (MADM)-based clonal analysis showed that while indeed all ET-PNs and IT-PNs derive from E12.5 multipotent RGPs, lineage progression occurs through fate restricted sub-lineages and in parallel. Most specifically, we found that the earliest neurogenic divisions of E12.5 RGPs generate very few individual PNs (direct neurogenesis). Instead, there neurogenic outputs are pure ET-PN and IT-PN sub-lineages. ET-PN lineages are always small and mostly composed of two PNs in the same DLs. IT-PN sub-lineages produce PNs in both DLs and ULs layers and include small self-consuming lineages and large translaminar (DL+ UL) subclones progressing through an IT-PN fate-restricted RGPs. This mode of simultaneous specification of sublineages through distinctive progenitors not amplifies the production of IT-PNs with respect to the number of ET-PNs, but also delays their birth-time relative to the birth of ET-PNs. Finally, we do not observe a stereotyped output from the divisions of multipotent RGPs at the individual level. It is not that every single division gives rise to an ET-PN sub-lineage on one side and an IT-PN sub-lineage on the other, but that the two daughter cells undertake on any fate or restriction (ET-PN, IT-PN, or none-i.e. remain multipotent) independently of the choice in the sibling cell. Moreover, we show that there is an equiprobable frequency of a daughter RGP to maintain the multipotent state of the parental RGP or to be specified as an IT-PN restricted RGP. Thus, at the individual level, there are multiple lineage trajectories for the lineage progression of a multipotent RGP. However, despite the lack of stereotyped patterns in the neurogenic divisions of multipotent RGPs, we show that the random specification of IT-PN fate-restricted RGPs generates a quantitatively and temporally stereotyped neurogenesis population-wise, and thus organizes cortical neurogenesis. Our findings provide a new perspective on the mechanisms generating PN diversity in the cerebral cortex and a novel framework for RGP lineage progression, with broad implications for neurodevelopmental disorders and evolution.

## MATERIALS AND METHODS

### Animals

Wild-type *C57BL6/J* mice (*Jackson Laboratories* (JAX#00064) were used for all experiments except for MADM. For MADM analysis we employed previously described transgenic mouse lines with MADM cassettes inserted in chromosome 11 (36): *MADM-11-GT* (*MADM^GT/GT^*; Jackson Laboratories JAX #013749) and *MADM-11-TG* (*MADM^TG/TG^*; Jackson Laboratories JAX #013751). We crossed *MADM-11-GT* mice with the driver line *Emx1*-CreERT^2^(37) for specific cortical expression. These mouse lines were kept in Mixed C57/Bl6, FVB, and CD1 genetic background.

Males and females of WT or the desired genotypes were used randomly for all experiments. For pregnant dams, the morning of the appearance of a vaginal plug was defined as embryonic day (E) 0.5. Animals were housed at 21±1°C ambient temperature and 40-55% humidity under 12:12-hour light/dark cycle. Water and food were provided *ad libitum.* Animals were bred and maintained following the guidelines from the European Union Council Directive (86/609/ European Economic Community). All procedures involving the handling and euthanasia of animals complied with European Commission guidelines (2010/63/EU). All animal procedures were approved by the CSIC, the Community of Madrid Ethics Committees on Animal Experimentation, and by the Austrian Federal Ministry of Science and Research, in compliance with national and European legislation (PROEX 123-17; 124-17; 158.7/21; 230.7/21; 159.5/22; BMWFW-66.018/0006-WF/V/3b/2017 and BMWFW-66.018/0015-WF/V/3b/2017).

### CTB injections for retrograde labeling

Retrograde labeling was performed by injecting cholera toxin subunit B (CTB) conjugated to Alexa Fluor 488, 555, or 647 (*Thermo Fisher Scientific*, ref. C-34775, C-34776, and C-34778) into the CC (33, 35) the IC, the CerbPed or the dorsal striatum. Stereotaxic coordinates (in millimeters) and CTB volumes are reported in Tables 1 and 2 shown below. For CTB injections performed between P0-P5, to ensure the completion of neuronal migration and NeuN expression, we sacrificed the animals at P10. For other injections, animals were sacrificed two days post-CTB injections to allow for retrograde transport of the tracer. In MADM experiments, two injections were performed into the CC using different coordinates along the anteroposterior axis to cover the entire tract width in the SS cortex. Anesthesia during the surgical procedure was obtained using isoflurane/oxygen (3% isofluorane for induction, 1.5-2% isofluorane for maintenance during the surgery). Anesthetized animals were placed on a stereotaxic frame (*Harvard Apparatus*) with a mouse neonatal adapter (*Stoelting*). CTB, diluted at 0.5% in phosphate-buffered saline (PBS), was injected with a Drummond Nanoject II Auto-Nanoliter Injector using 30 mm pulled glass micropipettes (*Drummond Scientific*, ref. 3000205 A and 3000203 G/X). CTB was delivered at a rate of 9.2nl per injection pulse with a maximum frequency of one pulse per second (9.2nl/s) to minimize damage until the desired total volume was achieved. Animals were perfused for analysis at least 48 h post-injection.

**TABLE 1:** CTB coordinates and volumes for CC injections.

| Injection site | Postnatal day (P) | Injected volume (nL) | Angle | Coordinates |  |  |
| --- | --- | --- | --- | --- | --- | --- |
|  |  |  |  | AP | ML | DV |
| **Somatosensory CC** | P1/P0 | 230 | 0 | -1 | +0.5 | -0.8 to -1.1 |
|  | P2 | 230 | 0 | -1 | +0.5 | -0.9 to -1.2 |
|  | P3 | 322 | 0 | -1 | +0.5 | -0.95 to -1.25 |
|  | P4 | 322 | 0 | -1 | +0.5 | -1.05 to -1.35 |
|  | P5 (anterior) | 460 | 0 | -1 | +0.5 | -1.05 to -1.35 |
|  | P5 (posterior) | 460 | 0 | -1.4 | +0.5 | -1.05 to -1.35 |
|  | P10 | 460 | 18 | -1.4 | +0.7 | -1.3 to -1.6 |
|  | P16 | 575 | 18 | -1.4 | +0.7 | -1.6 to -1.9 |
|  | P30 | 575 | 18 | -1.4 | +0.7 | -1.7 to -2 |

**TABLE 2:**
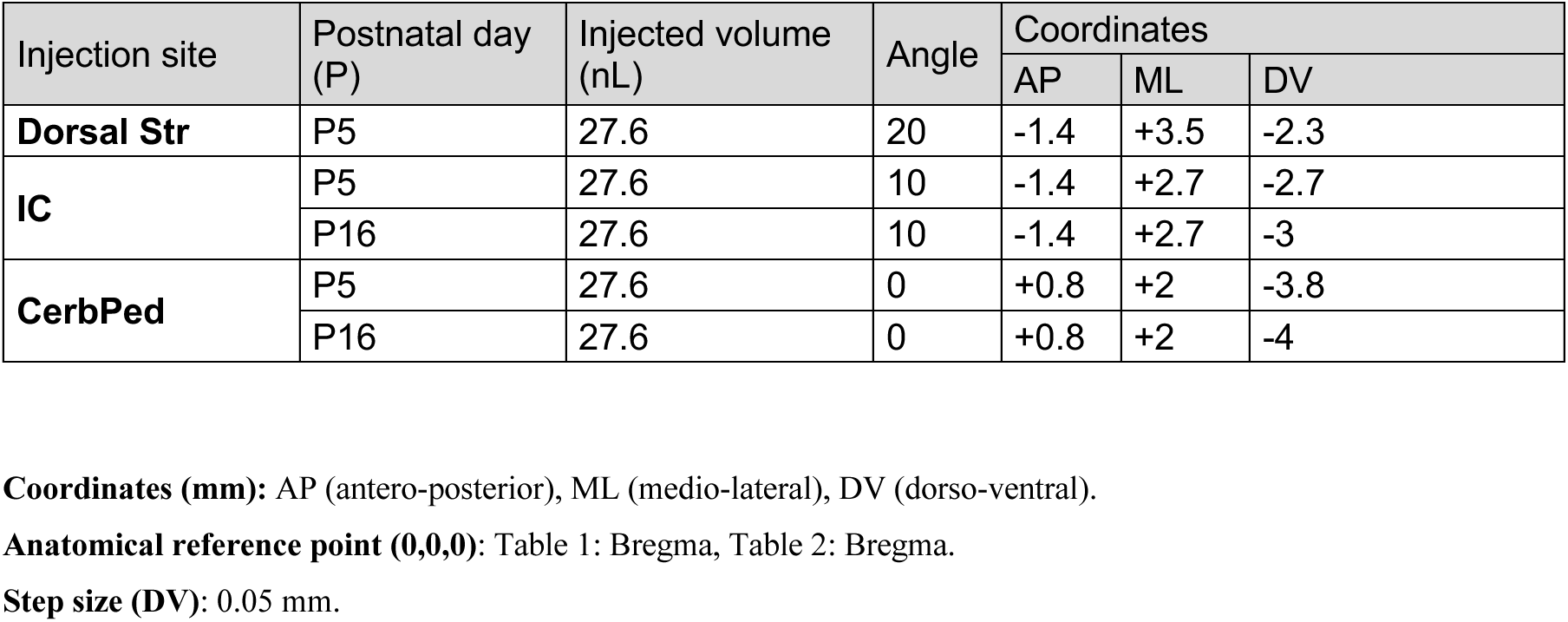
CTB coordinates and volumes for ET injections.

### EdU birth-dating

Pregnant dams were intraperitoneally injected with the thymidine analog, ethynyl deoxyuridine (EdU, 50 mg/kg body weight) using a 30G needle at different stages of embryonic development (E11.5, E12.5, E13.5, E14.5 or E15.5). The pups were cardially perfused at P10. Brains were collected, sectioned, and processed for the detection of EdU using the Click-It Alexa Fluor imaging kit according to the manufacturer’s instructions (*Thermo Fisher Scientific, ref C-10337 and C-10340*).

### MADM genetic clonal labeling

*MADM-11^GT/GT^; Emx1-CreER^T2+/−^*mice were crossed with *MADM-11^TG/TG^* mice using males and females of both strains indistinguishably. MADM clone induction was performed as previously described (8, 38). Timed pregnant dams were injected intraperitoneally at E12.5 or E13.5 with tamoxifen (TM) (*Sigma-Aldrich, ref. T5648-5G*) dissolved in corn oil (*Sigma-Aldrich, ref. C8267*) at a dose of 2 mg/pregnant dam (stock prepared at 20mg/mL). Embryos were recovered at E18–E19 through cesarean section, fostered and raised for CTB injection at P5, and sacrificed at P10 for analysis.

MADM-based clonal analysis using *MADM-11* transgenic lines has previously been validated to identify clusters of cells, each representing individual clonal units (9, 36, 38). The TM dose used in this study has been proven to achieve sparse labeling (8). A 300 mm minimum distance was used to determine separate clones in brains with multiple cell clusters.

### MADM taxonomy

To identify MADM clones, we first analyzed GFP^+^ and tdTomato^+^ cell clusters in each section sequentially, in the rostral to caudal order. High-magnification images of each individual neuron were acquired for assessing CTB^+^ labeling. Each clone was individually annotated for CTB labeling and layer occupancy in an atlas (see Clone Atlas in the Supplemental Information). For MADM taxonomy, we positively assigned IT-PN identity while we applied a negative classification for ET-PNs. Clones and subclones were categorized into three classes: IT-PN, consisting exclusively of IT-PNs (i.e. UL-PNs and CTB^+^ DL-PNs); ET-PN, consisting exclusively of CTB^-^ DL-PNs; and Mixed, containing both IT-PNs and ET-PNs. As a control, to ensure the correct discrimination of DL IT-PNs from ET-PNs in MADM experiments, we verified that CTB labeling efficiencies in MADM^+^ PNs were comparable to the values we obtained in WT mice in NeuN^+^ cells. Control experiments confirmed CTB labeling efficiency of MADM outputs comparable to WT animals (Figure **S5A**), and showed cell death rates and UL/DL-PN ratios (Figure **S5B** and S5C) consistent with previous reports (8).

### In utero electroporation (IUE) and plasmids

IUE was performed as previously described (39). Pregnant mice were anesthetized as described above. The uterine horns were extracted following a longitudinal incision in the epithelial tissue and laparotomy in the muscle. DNA solutions (1µg/µl CAG-*Gfp*, pCIG2-*Pou3f1* or CAG-*Pou3f2*, or a mix of shPou3f1 and shPou3f2, 0.8 µg/µl each) were prepared with 0.1% Fast Green in ddH2O (*Sigma-Aldrich, ref. F7252*). The plasmid solutions were injected in the lateral ventricle of the embryo using a 30 mm pulled glass micropipette. 5 pulses of 50 ms and 33, 36, 38V for E13.5, E14.5 and E15.5 respectively, were applied using external paddles (*Sonidel Limited, ref. CUY650P5*) oriented to target the S1 area, and an ECM 830 electroporator (BTX, ref. 650052). After, the uterine horns were replaced inside the abdominal cavity, and the incision was sutured. An intraperitoneal injection of carprofen solution (*IsoFlo. Zoetis Inc.*) was applied to prevent postoperative pain. For E13.5, a cesarean section at E19.5 was performed if required. Pups were allowed to develop normally until the stage of interest. The plasmids electroporated were: *CAG*-*Gfp* (Addgene, plasmid #11150), *pCIG2-Pou3f1* (gift from Michelle Cayouette), *CAG-Pou3f2* (Addgene, plasmid #19711). Immunostaining against GFP was performed as explained below for signal amplification.

### *In vitro* validation of short hairpin (sh) RNAs

To validate the efficacy of the shRNAs *in vitro,* N2A cells were cultured and transfected with the respective plasmids using Effectene (*Qiagen, ref. 301425*). Four shRNAs targeting Pou3f2 and five shRNA against Pou3f1 were tested, alongside a scrambled control. shRNAs were co-transfected with the overexpression plasmids (*CAG-Pou3f2* and *CAG-GFP*, or *pCIG-Pou3f1*). Transfection efficiency was monitored using GFP as a reporter. After 72 hours, total RNA was extracted from the cells using the RNAspin Mini Kit (*GE Heatlhcare*). RNA purity and concentration were verified using a NanoDrop spectrophotometer. cDNA synthesis was performed with 2 µg RNA using M-MLV reverse transcriptase (*Promega, ref. M1701*) according to the manufacturer’s protocol. Real-time PCR was conducted in 10 μl reaction volume using SYBR Green Master Mix (*Thermo Fisher Scientific, ref. A46012*). The QuantStudio 3 Detection system (*Applied Biosystems*) was employed under standard Applied Biosystems cycling parameters (40 cycles: 95°C for 15 seconds, 60°C for a 45 seconds). Primer sequences were designed with BioRender, and cDNA dilutions were optimized to match the amplification efficiencies of housekeeping genes (18S rRNA and Rpl13A). Relative mRNA expression was calculated following the ΔCt method, using the Best Keeper index (40) as a reference for the total mRNA load in the sample

### Perfusion and tissue collection

Mice were deeply anesthetized by intraperitoneal injection of Xylazine (*Xilagesic, Calier*) and Ketamine (*Imalgene Merial Laboratorios*) solution. Animals were transcardially perfused with PBS and either ice-cold 10% formalin solution (*Sigma-Aldrich, ref. HT501128-4L*), or 4% paraformaldehyde (*Sigma-Aldrich*, *ref. 441244-1KG*) in PBS for MADM experiments. Brains were dissected and post-fixed in the fixative for 4-12 h. Cryoprotection was achieved by immersion in 30% sucrose solution in PBS (*Merck, ref. S0389*) for 48 h. Then brains were embedded in Tissue-Tek O.C.T (*Sakura Tissue-Tek, ref.4583*) and stored at −20°C until further use. Tissue was sectioned with a cryostat (*Thermo Fisher Scientific*) in coronal orientation at 50 µm thickness, collected in PBS, stained when required, and mounted onto glass slides. Brain cryosections containing MADM clones were collected and kept in the same left-right orientation and in serial order, in multiple 24-well plates containing PBS.

### Brain slice processing, immunostaining, and antibodies

Free-floating sections were used for immunostaining procedures. Sections were immunostained with antibodies overnight at 4°C or room temperature (RT) in a solution composed of PBS with 0.5% Triton (PBS-T) (*Sigma-Aldrich, Triton^TM^ X-100*) and 5% Fetal Bovine Serum (*FBS, Thermo Fisher Scientific, ref: A5256701*), and after washed and incubated with the secondary antibodies diluted in 2% FBS in PBS-T. Cell nuclei were stained by 15 min incubation with PBS containing 4’,6-diamidino-2-phenylindole (DAPI) (1:1000 dilution*, Thermo Fisher Scientific*, *ref: D1306*). Sections were embedded in aqua polymount (*Aqua-Poly/mount. Polysciences, Inc. ref: 18606*) and stored at 4°C until image acquisition. For MADM analysis, brain sections were mounted on Superfrost glass slides (*Thermo Fisher Scientific*) and embedded in mounting medium containing Mowiol 4-88 (Carl Roth, *ref: Art.-No. 0713*) and 1,4-diazabicyclooctane (DABCO; Carl Roth, *ref: Art. No. 0718*). For embryonic tissue, brain sections were mounted on Superfrost glass slides (*Thermo Fisher Scientific*) and boiled for antigen retrieval. Sections were immersed in a citrate solution (*Sigma Aldrich*, ref: C9999) and heated to 110° for one minute in a pressure cooker (*Biocare Medical*, ref: DC2012).

Primary antibodies: mouse monoclonal anti-NeuN (1:500, Merck Millipore, ref: AB377/A60), rabbit polyclonal anti-GFP (1:500, Thermo Fisher Scientific, ref: A-11122), mouse monoclonal anti-Tbr2 (1:100, eBioscience-Thermo Fisher Scientific, 14-4875-82/Dan11mag), rabbit polyclonal anti-Pou3f1 (1:100, Abcam, ref: ab272925), rabbit monoclonal anti-Ctip2 (1:250, Abcam, ref: ab240636). Secondary antibodies: goat polyclonal anti-mouse 488/546/647 (1:500, Thermo Fisher Scientific, A-11006/A-11030/A-21236), goat polyclonal anti-rabbit 488/546/647 (1:500, Thermo Fisher Scientific, A-11034/A-11035/A-21245).

### Image acquisition

Confocal image acquisition was performed using inverted confocal microscopes LSM 800 series (Zeiss), SP8, or Stellaris5 (Leica). Optical z-sections were obtained by taking 1-3.5μm serial confocal images with LAS AF v1.8 software (Leica) or ZEN Blue 3.8 (Zeiss). Mosaic images were obtained with the 20x/0.8 objective at 512×512 or 1024×1024 resolution (8 bits). High-magnification images were acquired with the 63x/1.4 oil immersion objective at 1024×1024 resolution (8 bits). All images were analyzed using Fiji-ImageJ (71) and ZEN Blue 3.8 (Zeiss). For MADM clones, sections were first screened using an axioscope (Zeiss Axio Imager, Zeiss) coupled to a CoolLED p300 SB light source (CoolLED) and equipped with 10x/0.45 and 20x/0.8 objectives (Zeiss). Green and red fluorescence were observed using an HC-dualband GFP/DsRed filter (F56-420, AHF).

### Quantification of IUE and EdU, CTB-labeled populations

To delimit S1-Bf equivalent areas in the developing brain, we used the Developing Mouse Reference Atlas from Allen Brain Atlas (http://atlas.brain-map.org/) and anatomical landmarks. For EdU, IUE, and injections in the IC, CerbPed, or dorsal striatum experiments, the entire S1Bf region was quantified after the area and cortical layers were outlined based on anatomical landmarks and their distinct cell densities. To quantify CTB^+^ cells, we analyzed confocal plane z-stack images spanning the width of each cell of interest (33, 35). Cells are marked as positive if the CTB fluorescent signal surrounds the DAPI nuclei of the analyzed cell (NeuN^+^, EdU^+^, GFP^+^). For CTB^+^NeuN^+^ quantification after injections in the CC, 250 (50 per layer) NeuN^+^ cells were randomly selected within an ROI depicting a column in the S1Bf on the hemisphere contralateral to the injection site. Data are provided as the percentage of CTB^+^ neurons out of NeuN^+^ cells. A minimum of two sections (technical replicates) were analyzed per individual brain. At least three animals (biological replicates) were analyzed per condition/stage.

### Analysis of single-cell RNA seq dataset

To cluster RGPs and study differential gene expression over developmental time, we analyzed the single-cell RNA sequencing atlas of the developing neocortex published by Di Bella et al 2021. We selected cortical cells from E12.5 to E15.5 samples. The analysis included the following steps using *R software* (DOI: 10.5281/zenodo.14609057): (i) **Normalization and Filtering:** Counts were standardized for each time point trough the normalization of the feature expression measurements for each cell by the total expression, scaling by a factor of 10,000, and applying a log+1 transformation. The dataset was filtered to include only samples from embryonic stages E12 to E15. Prior to this analysis, the data was already pre-filtered to retain high-quality cells, with mitochondrial reads <7.5% and detected genes >500. (ii) **Feature Selection and Scaling:** We identified the 3000 most variable features using the FindVariableFeatures function with the ‘vst’ selection method. During scaling, we regressed out variability associated with ‘nCount_RNA’ and ‘nFeature_RNA’ to correct for technical artifacts related to RNA capture and sequencing depth. (iii) **Dimensionality Reduction and Clustering:** we performed linear dimensionality reduction using Principal Component Analysis (PCA) on the scaled data, retaining 50 principal components. A k-nearest-neighbors graph was constructed based on Euclidean distance in PCA space using these 50 components (FindNeighbors, dims = 1:50). Cells were clustered using the Louvain algorithm implemented in Seurat, with a resolution parameter set to 0.7 (FindClusters). (iv) **Visualization:** For visualization, Uniform Manifold Approximation and Projection (UMAP) was applied to the first 15 principal components (RunUMAP). (V) **Cell Annotations and Differential Expression:** Cell annotations for embryonic stage and cell type were incorporated into the analysis. Differentially expressed genes were identified using FindAllMarkers in Seurat with the Wilcoxon Rank Sum test, applying a Bonferroni correction for multiple testing (adjusted p-value < 0.05). Only genes detected in at least 25% of the cells within a cluster and showing a minimum 0.25-fold difference (log-scale) between the cells in the cluster and all remaining cells were considered.

### Statistical analysis

All experimental conditions include a minimum of 3 mice and two coronal sections per brain, except for MADM experiments in which we analyzed the entire brain. Only the animals in which CTB labeling was not efficient or in which IUE experiments targeted other cortical areas than the somatosensory cortex were excluded from the analysis. Results show the sample mean +/- the standard error of the mean (SEM) or the median. The statistical groups were determined by embryonic and/or postnatal experimental day, treatment, or layer. Before applying parametric statistical analysis, sample normality was tested using the D’Agostino-Pearson or Sapiro-Wilk test. Data visualization was performed using GraphPad Prism 10.0 (GraphPad Software). The results were compared using different tests depending on the study object and the number of variables. The student t-test was employed when comparing the mean values of one variable between two groups; one- or two-way analysis of variance (ANOVA), followed by a posthoc test, Dunnett’s, Tukey’s, or Šídák’s multiple comparisons test as specified in the figure legends. Kruskal Wallis (KW) test or Wilcoxon matched-pairs tests were used for samples that did not follow a normal distribution. Fisher exact test and Chi-square test were applied to compare distributions. Linear regression and correlation (Spearman test) analysis were used to evaluate the relationship between variables. Statistically significant differences were established at p <0.05 (*), p < 0.01 (**), p < 0.001 (***), and p <0.0001 (****), “n.s.” stands for non-significant.

Chi-square and permutation tests were combined to assess the significance of the association between clone/subclone type (IT-PN only, ET-PN only, Mixed) and the type of division of the clone/subclone (Proliferative, Asymmetric, Terminal). A Chi-square statistic was calculated for the observed data and then compared to chance using the R function “chisq.test”. The “type of clone/subclone” column was permuted 1000 times, and a chi-square statistic was calculated for each permutated dataset. The p-value was calculated as the proportion of permuted chi-square statistics that are greater than or equal to the observed chi-square statistic.

Estimates of death rate in MADM clones and of the maximal potential frequency of IT-PN clones emerging from the death of ET-PNs in a Mixed clone: The cell death rate (probability=0.08/cell) was calculated based on the frequency of orphan/single-color subclones, as previously described (8). This value aligns with earlier reports (8, 22). For the estimate of misclassification errors due to cell death, we considered that there could be “ghost”-Mixed clone/lineage misclassified as pure clone/lineage due to the cell death of ET-PNs or IT-PNs. The maximum frequency of a ghost-mixed clone/lineage was calculated considering that, in a Mixed lineage, the frequency of losing ET-PN or IT-PN branches cannot exceed the progenitor death rate. Thus, we calculated a maximum error of our measurements as the probability of the death of either the ET-PN or IT-PN progenitor in a mixed clone or lineage. For example, the frequency of mixed clones in the population labeled at E12.5 is 0.78. Thus, the maximum frequency of death of ET-PN or IT-PN precursor in a mixed clone/lineage = (frequency of mixed clones/lineage) x 0.08 (death rate of a subclone) x 0.5 (probability of ET-PN or IT-PN). The errors we obtained exclude statistically the possibility that the frequencies of pure clones and lineages we observed are due to cell death. Examples:

- **Error in the measurement of E12.5 ET-PN clones** Maximum frequency of death of ET-PN precursor in a mixed clone/lineage = (frequency of mixed clones/lineage) x 0.08 (death rate of a subclone) x 0.5 (probability of ET-PN) = 0.03 (3%). Which provides a final frequency of **10,6 ±3 %.**
- **Error in the measurements of E12.5 ET-PN lineages** Maximum frequency of death of ET-PN precursor in lineage = = 0.48 (frequency of mixed lineages) x 0.08 x 0.5 = 0.02 (2%). Which provides a final frequency of **24** ±2 %.
- **Error in the measurements of E12.5 IT-PN lineages** Maximum frequency of death of IT-PN precursor in a mixed clone/lineage = (frequency of mixed clones/lineage) x 0.08 (death rate of a subclone) x 0.5 (probability of IT-PN) = 0.03 (3%). The experimental frequency of E12.5 IT-PN clones is 12%, which, with the error accounting for cell death, provides a final frequency of **12 ±3 %.**

Alternative statistical calculations taking into account additional considerations provide even lower error rates due to cell death. For instance, by considering the size of the clone/lineage, we can refine our estimates: *e.g*., in an IT-PN clone of 6 neurons, the error due to the death of an ET-PN in a mixed clone of 7 neurons can be calculated using the experimental frequency of an IT-PN clone with a size of 7 and the probability of the dying cell being an ET-PN (1/8). This yields an error frequency for clones/lineages of 8 neurons of 0.000196 (0.0196%).

## RESULTS

### Developmental callosal projections as a specific marker for IT-PNs

To determine whether early callosal projections specifically characterize IT-PNs, we investigated the interhemispheric projections of DL-PNs using injections of retrograde tracer molecules (cholera toxin subunit B [CTB] fluorescent conjugates) into the CC (Figures **1A-C**). Unlike cortical plate injections, this approach saturates the callosal tract and enables quantification of the absolute numbers of PNs with developmental or mature callosal axons. A comprehensive description of the methodology, including demonstrations of specificity, saturation, and efficacy, has been previously reported (33).

**Figure 1.**
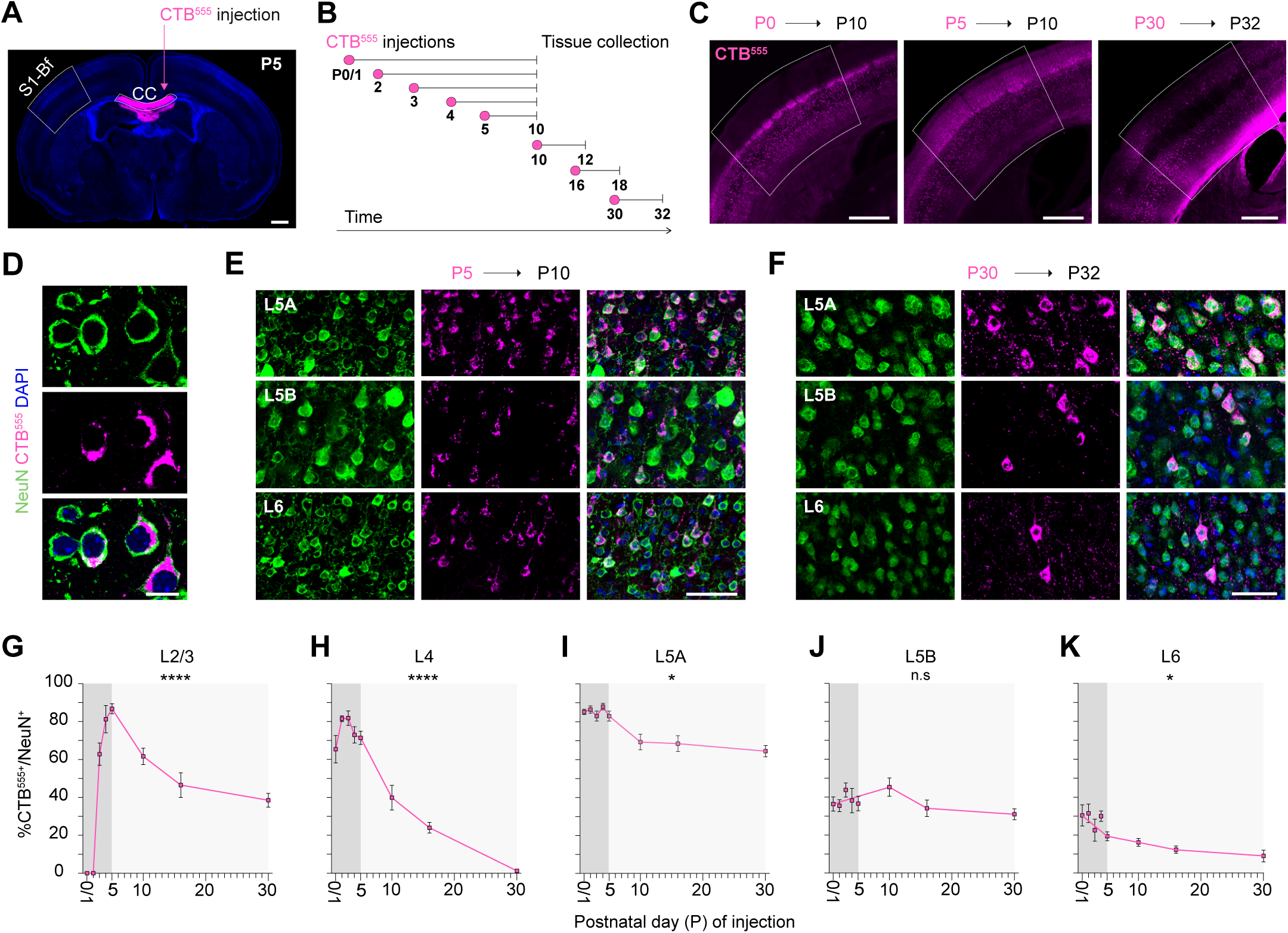
PNs across all cortical layers extend developmental callosal axons. (**A**) CTB-555 signal confined within the CC two hours post-injection. (**B**) Experimental paradigm for CPN analysis: saturating CTB injections into the CC were performed (i) at P0/1, P2, P3, P4, or P5 and brains analyzed at P10, once neuronal migration is completed, or (ii) at P10, P16, or P30, and brains analyzed 48 hours after the injection. (**C**) CTB-555 signal in the S1-Bf cortex contralateral to CC injection sites at P0 (left), P5 (middle), and P30 (right). (**D**) High-magnification image of L6 PNs labeled for NeuN (green), CTB-555 (magenta), and DAPI (blue) from P5-injected brain analyzed at P10. (**E** and **F**) NeuN (green), CTB-555 (magenta), and DAPI (blue) signals in the DLs of brains injected and analyzed at the indicated times. (**G-K**) Layer-specific dynamics of callosal axonal remodeling. Animals were injected and analyzed as in (B). The graph represents the percentages of CPNs (CTB^+^NeuN^+^) per cortical layer (y-axe) and time of injection (x-axe). Mean ± SEM (*n* = 500 cells, *n* = 2 sections/animal; *n* ≥ 4 animals/stage). One-way ANOVA (CTB^+^NeuN^+^ vs. Time) : (G) **** p-value <0.0001, (**H**) **** p-value <0.0001, (I) * p-value =0.0223, (J) p-value =0.1399 (n.s), (K) * p-value =0.038. **Scale bars:** 500 μm (A and B), 10 μm (D), 50 μm (E and F),

To examine axonal refinement throughout development, we injected animals at ages ranging from postnatal day (P) 0 to P30 (Figures **1A-C**). To avoid confounding effects from other cell types, we immunolabeled for NeuN and quantified the proportion of PNs with callosal axons (CTB^+^NeuN^+^ cells) (Figures **1D-F** and **S1A** and **S1B**). Analysis in the primary somatosensory cortex (S1) revealed that callosal projections peak between P0 and P5 (Figures **1C-K**, **S1A-G**), confirming the common callosal projection neuron (CPN) identity of early UL-PNs (Figures **1G** and **1H**, **S1A-G**) (33, 41, 42).

In the DLs, our experiments demonstrated that L5A is principally composed of CPNs, which, unlike UL-PNs, undergo minimal pruning from birth to adulthood (Figure **1I**). Additionally, we found that approximately 30% of the PNs in L6 and L5B display early callosal projections, revealing their early CPN identity (Figures **1J** and **1K**). While the number of L5B CPNs remained stable from P0 to P30, pruning reduced the proportion of L6 CPNs to about 10% in adults (Figures **1J** and **1K**). Furthermore, consistent with the projection patterns of adult IT-PNs, many DL-PNs with callosal axons–preferentially from L5A and L5B– projected to the dorsal striatum, both inter-hemispherically and intra-hemispherically (Figure **S2A-G**) (43).

We next investigated whether DL-PNs with developmental callosal axons might express a hybrid PN identity and also extend projections to ET territories. To address this, we conducted a series of dual-labeling experiments by combining injections into the CC with subsequent injections into either the internal capsule (IC) or cerebral peduncle (CerbPed). These two anatomical sites were targeted as they harbor the major ET projection routes, including corticothalamic, corticopontine, and corticospinal fibers (44) (Figure **2A**). One group of animals received CC injections at postnatal day 3 (P3) followed by IC or CerbPed injections at P5, while another group received both injections at P16 to assess a more advanced stage of axonal refinement (Figure **2B**).

**Figure 2.**
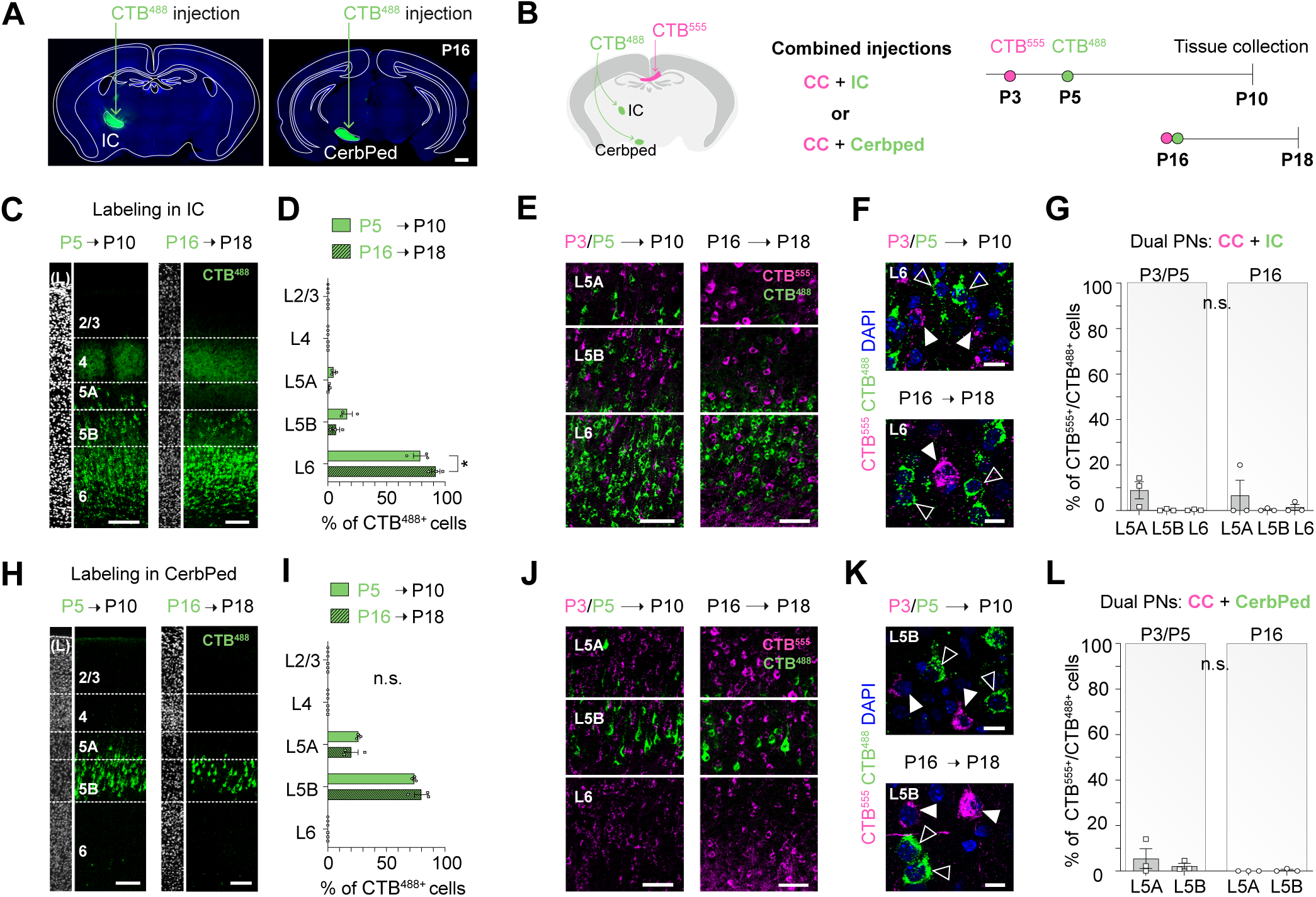
Early postnatal axonal tracing from the CC differentiates IT-PNs from ET-PNs. (**A**) CTB-488 signal confined within the IC (Left) or the CerbPed (Right) two hours post-injection. (**B**) Injection scheme: sequential (P3 and P5) or simultaneous (P16) injections into the CC (magenta) and the IC or the CerbPed (green) to analyze dual-projection neurons. (**C** and **H**) CTB labeled S1-Bf PNs projecting through the IC (C) or the CerbPed (H) following injections as in (A). (**D** and **I**) Percentage of CTB-488^+^ cells (ET injection) per cortical layer after injections as in (B). Mean ± SEM (*n* = CTB-488^+^ cells in the S1-Bf, *n* = 2 sections/animal; *n* = 3 animals/group). Two-way ANOVA with Šidak’s posthoc test: * p -value _L6 P5 vs L6 P16_ =0.0147 in (D). (**E** and **J**) CTB-488 (green) and CTB-555 (magenta) signals segregated into different PNs after injections as in (B). (**F** and **K**) High-magnification images from (E and J) CTB-488^+^ (green, empty arrowheads) and CTB-555^+^ cells (magenta, white arrowheads). (**G** and **L**) Percentage of CTB-555^+^ PNs over CTB-488^+^ PNs after IC (G) or CerbPed (L) injections as in (B). Mean ± SEM (*n* = CTB-488^+^/layer, *n* = 2 sections/animal; *n* = 3 animals/group). The two-way ANOVA analysis for comparison with a theoretical zero showed no significant main effects. **Scale bars:** 500 μm (A), 100 μm (C and H), 50 μm (E and J), 10 μm (F and K).

Both types of ET injections demonstrated specificity and labeled cells exclusively in the DLs (Figures **2C**, **2D**, **2H** and **2I**). The distribution of labeled cells across layers remained largely consistent between P5 and P16 injections, indicating limited pruning during this period (Figures **2C**, **2D**, **2H** and **2I**). Notably, the vast majority of labeled cells were single-positive, i.e., labeled exclusively by either the callosal or ET injection (Figures **2E**, **2F**, **2J**, **2K**, **2G** and **2L**). The absence of dually labeled cells indicated that ET-PNs do not project inter-hemispherically, nor in adults or during development, confirming that developmental callosal axons are a hallmark of IT-PNs. These experiments demonstrated that early callosal projections are a reliable indicator of intra-telencephalic (IT) PN identity, not only for UL-PNs but also for DL-PNs.

### Sequential production of Extra- and Intra-telencephalic projection neurons

Retrograde tracing from the CC at early postnatal stages specifically distinguishes IT-PNs from ET-PNs and serves to discriminate the two types in the DLs. To elucidate the birth times of these PN subtypes, we administered single doses of 5-ethynyl-2’-deoxyuridine (EdU) to pregnant dams at embryonic days (E) 11.5, 12.5, 13.5, 14.5, or 15.5. In the first set of experiments, IT-PNs across all layers were subsequently identified by injecting cholera toxin subunit B (CTB) into the CC, and ET-PNs were identified with injections into the IC or the CerbPed of pups at postnatal day 5 (P5), a stage preceding major axonal refinement (Figure **1G-K**). We then quantified the distribution of EdU^+^ cells (newborn cells) and double-labeled CTB^+^EdU^+^ cells (newborn IT-PNs or ET-PNs) across all layers in the primary somatosensory cortex (S1) at P10 (Figures **3A-H** and **4A-G**).

**Figure 3.**
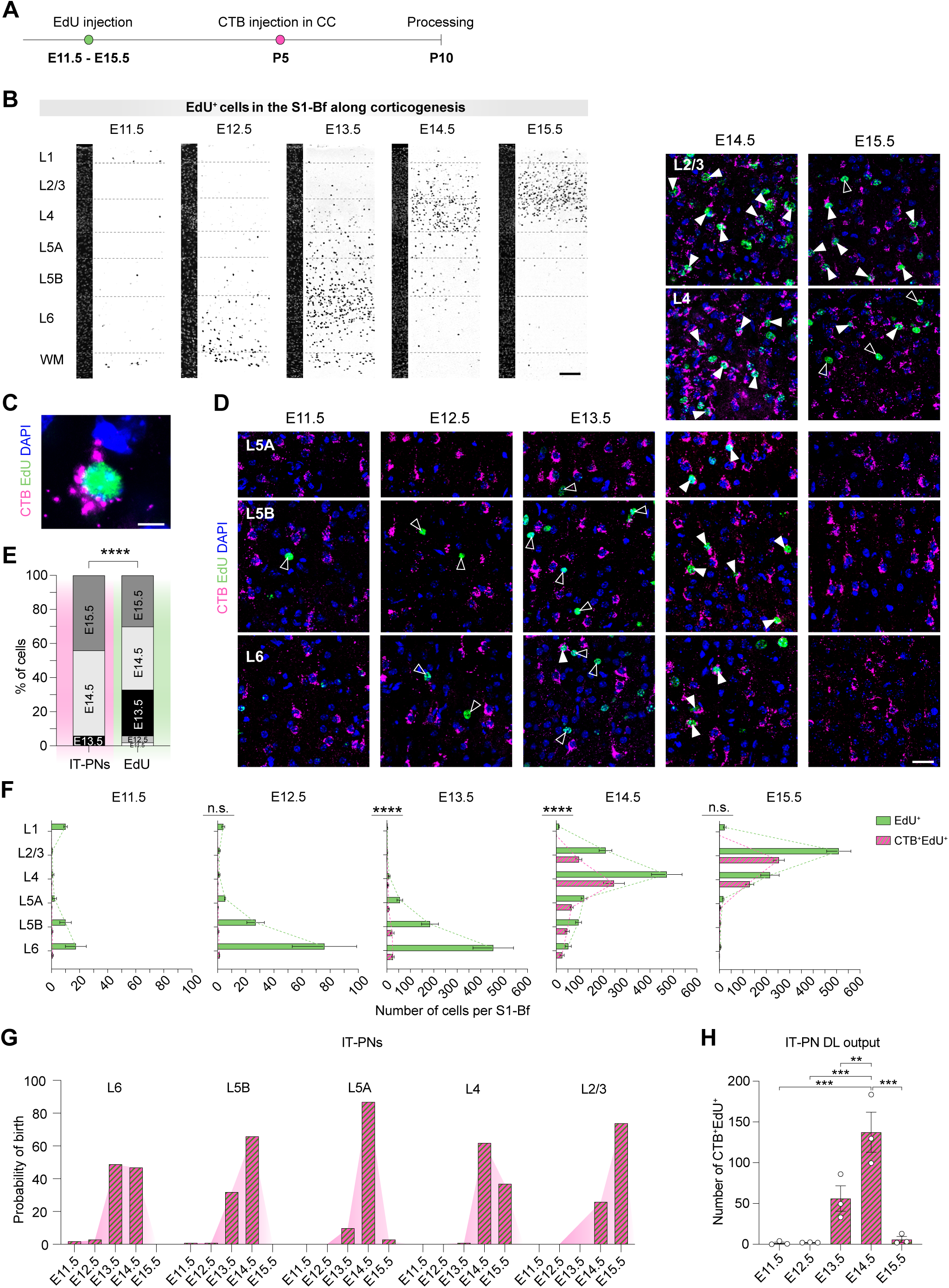
IT-PNs are born during a protracted period from mid neurogenesis. (**A**) Experimental design: EdU was injected at E11.5, E12.5, E13.5, E14.5, or E15.5, CTB-555 was injected into the CC at P5, and brains were analyzed at P10. (**B**) EdU labeling (gray) in the S1-Bf at P10 following EdU injections at the indicated embryonic stages. Dashed lines outline cortical layers. (**C**) High-magnification image of an IT-PN. CTB-555 (magenta), EdU (green) and DAPI (blue). (**D**) CTB-555 (magenta), EdU (green), and DAPI (blue) signals in the S1-Bf DLs (E11.5 – E13.5) or in both, the S1-Bf ULs and DLs (E14.5 and E15.5). EdU^+^ cells are indicated by empty arrowheads, CTB^+^ EdU^+^ cells by white arrowheads. (**E**) Fractions of CTB^+^ EdU^+^ (IT-PNs) and EdU^+^ cells born at the indicated stages. Data were calculated using the averages of the total number of IT-PNs or EdU^+^ cells per embryonic stage: (*n* = 2 sections/animal, *n* = 3 animals/stage). Chi-square test: **** p-value _IT-PNs vs EdU_ <0.0001. (**F**) Layer distributions of birthdated total cells (green) and IT-PNs (magenta) in the S1-Bf. Mean ± SEM (*n* = EdU^+^ or CTB^+^EdU^+^ cells/S1-Bf, *n* = 2 sections/animal; *n* = 3 animals/stage). Two-way ANOVA (CTB^+^EdU^+^ across consecutive stages) : (n.s) p-value _E11.5 vs. E12.5_ =0.4192, **** p-value _E12.5 vs. E13.5_ <0.0001, **** p-value _E13.5 vs. E14.5_ <0.0001, (n.s) p-value _E14.5 vs. E15.5_ =0.1174. (**G**) Proportion of IT-PNs per layer born at the embryonic stages in the x-axe. Data were calculated using the averages of the total number of IT-PNs per embryonic stage: (*n* = IT-PN number, *n* = 2 sections/animal, *n* = 3 animals/embryonic stage). (**H**) Absolute number of DL-IT-PNs per S1-Bf born at the indicated stages. Mean ± SEM (*n* = CTB^+^EdU^+^ cells/S1-Bf, *n* = 2 sections/animal; *n* = 3 animals/stage). One-way ANOVA with Tukey’s posthoc. *** p-value _E11.5 vs E14.5_ =0.0002, *** p-value _E12.5 vs E14.5_ =0.0002, ** p-value _E13.5 vs E14.5_ =0.0094, *** p-value _E14.5 vs E15.5_ =0.0003. **Scale bars:** 100 μm (B), 5 μm (C), 20 μm (D).

EdU birth-dating profiles aligned with classical studies, demonstrating smaller neuronal outputs at E11.5 or E12.5 than after E13.5 (Figure **3B**; **S3A-B**). Notably, the birth of IT-PNs (CTB^+^EdU^+^) (Figures **3C** and **3D**) exhibited a significant delay relative to the overall EdU^+^ output (Figures **3E**; **S3C**). As a control, immunolabeling for NeuN confirmed that CTB^+^EdU^+^ cells specifically identified PNs (Figure **S3C-E**). The majority of IT-PNs, including a substantial fraction of DL-IT-PNs, were labeled following E14.5 and E15.5 EdU injections. Only a small fraction of IT-PNs were labeled at E13.5 (Figure **3E**). More specifically, the data showed that half of L6 IT-PNs are born at E13.5 and half at E14.5; most layer L5B IT-PNs are born at E14.5; and the birth of layer L5A IT-PNs is tightly confined to E14.5. UL-PNs were labeled from E14.5 onwards, as expected (12, 45, 46) (Figures **3F-G**; **S3F** and **S3G**).

A parallel analysis of the birth times of ET-PNs showed that it is sharply restricted to E13.5 (Figures **4A-G**). Of note, E12.5 outputs were not labeled from CTB injections in the CC, nor from ET injections, indicating that these PNs represented subpopulations that do not extend long subcortical projections, such as the Tle4^+^ layer 6 PN subpopulation (28, 31).

**Figure 4.**
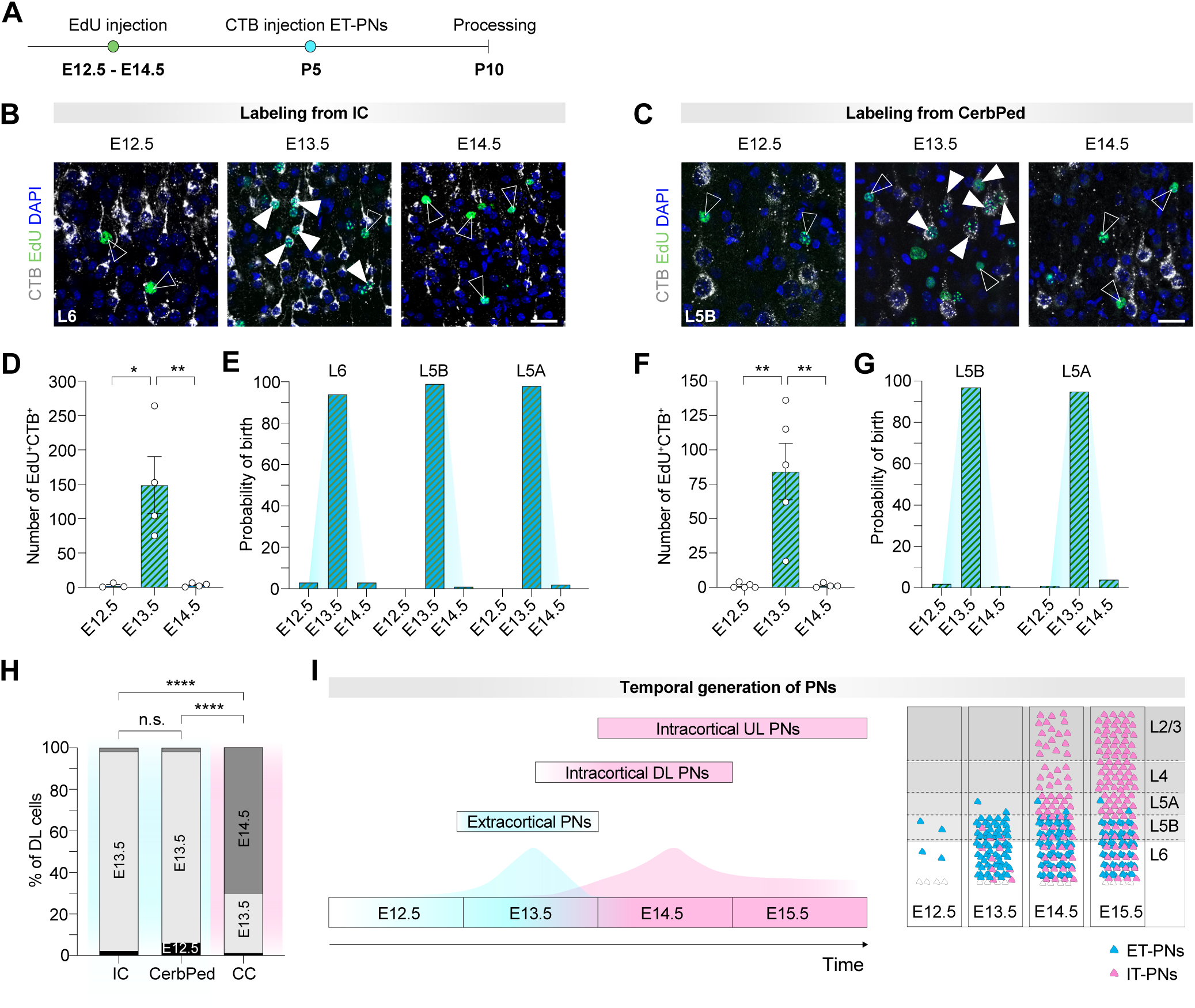
ET-PN production during a neurogenic burst at E13.5. (**A**) Experimental design: EdU was injected at E12.5, E13.5 or E14.5, CTB was injected into the IC or the CerbPed at P5, and brains were analyzed at P10. (**B** and **C**) Representative images of L6 (B) or L5B (C) PNs showing CTB (white), EdU (green), and DAPI (blue) labeling after IC (B) and CerbPed (C) injections. EdU^+^ cells are indicated by empty arrowheads, CTB^+^EdU^+^ cells by white arrowheads. Scale bar: 20 μm. (**D** and **F)** Total CTB^+^EdU^+^ cells per S1-Bf ROI after IC (D) or CerbPed (F) injections. Mean ± SEM *(n* = CTB^+^ EdU^+^ cells/S1-Bf*, n* = 2 sections/animal; *n* ≥ 3 animals/stage). One-way ANOVA with Tukey’s posthoc: (D) * p-value _E12.5 vs. E13.5_ =0.0137, (D) ** p-value _E13.5 vs. E14.5_ =0.0093, (F) ** p-value _E12.5 vs. E13.5_ =0.0016, (F) ** p-value _E13.5 vs. E14.5_ =0.0025. (**E** and **G**) Proportion of ET-PNs in the S1-Bf labeled from the IC (**E**) or the CerbPed (**G**) and born at the embryonic stages in the x-axe. Data were calculated with the averages of the total number of ET-PNs per embryonic stage *(n* = CTB^+^ EdU^+^ cells/S1-Bf*, n* = 2 sections/animal; *n* ≥ 3 animals). (**H**) Differences in the birthdates of ET-PNs and IT-PNs from the DLs. Data were calculated with the averages of the total number of DL-PNs labeled from the IC, CerbPed or CC per embryonic stage *(n* = DL-IT-PN / ET-PN number*, n* = 2 sections/animal; *n* ≥ 3 animals/embryonic stage/type of injection). Chi-square test: **** p-value _IC ET-PNs vs DL-IT-PNs_ <0.0001, **** p-value _CerbPed ET-PNs vs DL-IT-PNs_ <0.0001. (**I**) Schematic illustrating the neurogenesis of IT-PNs and ET-PNs in the S1-Bf.

To further analyze the sequential outputs from cortical progenitors, we performed in utero electroporation (IUE) experiments targeting progenitors at E13.5, E14.5, and E15.5 (**S4A-D**). Confirming the narrow window for ET-PN production, we observed green fluorescent protein (GFP)^+^ projections in the IC and thalamus exclusively in E13.5 outputs (Figure **S4E-F**), whereas in animals electroporated at E14.5 and E15.5, we found GFP+ projections in the CC and the striatum (Figure **S4G-J**).

Together, the experiments demonstrated that a neurogenic burst of ET-PNs at E13.5 precedes the birth of most IT-PNs (Figure **3H** and **4H-I**). Hence, IT-PNs and ET-PNs occupying the same DLs have completely different birthdates. This suggested that neurogenesis progresses hierarchically through the specification of projection subtypes in the progenitors rather than through the sequential generation of layer identities.

### IT-PN fate-restricted RGPs emerge from E12.5 multipotent RGPs

Our analysis of IT-PN and ET-PN production indicated that PN subtype specification is a hierarchical primary neurogenic decision and suggested a sequential origin within progenitors’ lineage progression. To investigate how individual RGPs specify and produce IT-PNs and ET-PNs, we combined our CTB injection protocol with MADM clonal analysis (6, 36) (Figure **S5A**). We used the MADM-11^GT/TG^; *Emx1-CreERT^2^* line to induce sparse labeling of dividing VZ progenitors following TM administration at E12.5 or E13.5. For IT-PN identification, we performed saturating injections of CTB into the CC at P5. We then individually assessed CTB labeling in each MADM-labeled cell at P10 (Figure **5A-C**). All UL-PNs were classified as IT-PNs, while DL-PNs were distinguished as IT-PNs (CTB^+^) or ET-PNs (CTB^-^) (Figure **5C**, **S6A** and raw MADM supplemental dataset provided as an annotated atlas). Based on subclone sizes, we inferred the division pattern of individual progenitors classifying MADM clones as proliferative (two subclones ≥4 PNs), asymmetric (majority subclone ≥ 4 PNs and minority subclone ≤3 PNs), and terminal (two subclones of ≤ 3 PNs) (Figure **S5B-H**). Proliferative and asymmetric divisions identify RGP divisions, while terminal divisions may not. We analyzed 75 clones labeled by TM inductions at E12.5 and 33 at E13.5.

**Figure 5.**
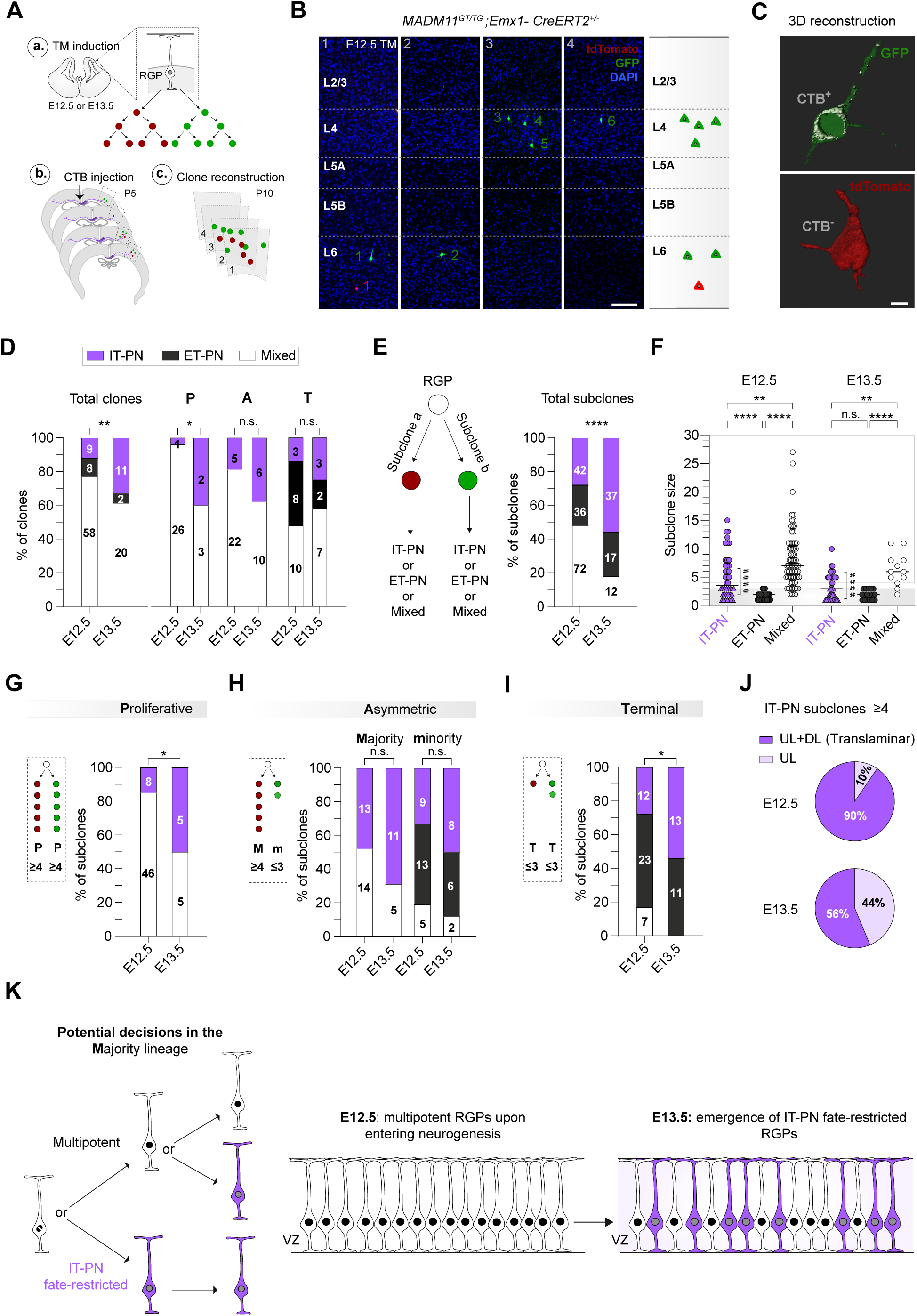
IT-PN fate-restricted RGPs emerge from E12.5 MADM-labeled divisions. (**A**) Experimental design: a. TM induction at E12.5 or E13.5, b. Saturating CTB injection into the CC at P5, c. Tissue processing and clone reconstruction at P10. (**B**) Consecutive brain sections of an asymmetric G2-X clone (panels 1-4) composed of a red (tdTomato) and a green (GFP) subclones. PN numbers/subclone in corresponding colors. Right-most panel: schematic of the reconstructed clone. (**C**) 3D Imaris reconstruction of a CTB^+^ IT-PN (GFP in green, CTB in white) and a CTB^-^ ET-PN (tdTomato in red). (**D**) Proportion of IT-PN, ET-PN and Mixed clones at E12.5 and E13.5. Chi-square test (IT-PN clones at E12.5 vs. E13.5): (**Total clones**) ** p-value =0.0086, (-**P**roliferative) * p-value =0.0105, (-**A**symmetric) p-value=0.1679, (-**T**erminal) p-value =0.4047. Bold numbers represent the number of clones per category. (**E**) (Left) Schematic of subclone definition and classification. (Right) Proportion of IT-PN, ET-PN and Mixed subclones at E12.5 and E13.5. Chi-square test: **** p-value <0.0001. Bold numbers represent the number of clones. (**F**) Subclone sizes after E12.5 (n=150 subclones) or E13.5 (n=66 subclones) TM induction. Data represent the median. Klustal-Wallis test with Dunn’s posthoc: (E12.5) **** p-value _IT vs ET_ <0.0001, ** p-value _IT vs Mixed_ =0.022, **** p-value _ET vs Mixed_ <0.0001, (E13.5) (n.s.) p-value _IT vs ET_ =0.2066, ** p-value _IT vs. Mixed_ =0.0049, **** p-value _ET vs Mixed_ <0.0001. Kolmogorov-Smirnov test comparing IT-PN sizes (small (≤3) vs. large (≥4)): ^####^ p-value <0.0001. (**G-I**) Proportion of IT-PN, ET-PN and Mixed subclones across proliferative (G), asymmetric (H), or terminal (I) divisions. Chi-square test: (G) * p-value = 0.0111, (H) (M, left) (n.s.) p-value=0.1885, (m, right) (n.s.) p-value=0.5523, (I) * p-value = 0.0321. Bold numbers represent the number of subclones. (**J**) Fraction of translaminar or pure-UL, IT-PN sub-lineages in proliferative and majority subclones. (**K**) Schematic summarizing key findings: Multipotent RGPs generate IT-PN fate-restricted and multipotent sub-lineages. The proportion of IT-PN RGPs increases between E12.5 and E13.5, gradually directing neurogenesis towards IT-PNs. (D-I) Clone/subclone types: IT-PN (purple), ET-PN (black) and Mixed (white). **Scale bars:** 100 μm (B), 5 μm (C).

77% of the clones produced from E12.5 MADM divisions include both IT-PNs and ET-PNs (Mixed clones). Specifically, 96% of the proliferative clones and 81% of the asymmetric clones were Mixed clones (Figure **5D**). Hence, overall and in agreement with previous studies, the E12.5 RGP population is multipotent (8, 10). From here on we refer to multipotent RGPs as those RGPs producing both ET-PNs and IT-PNs. 33% of E12.5 clones were pure ET-PN (11%) or IT-PN (12%) lineages. Notably, pure ET-PN clones were exclusively produced from terminal divisions (Figure **5D**). They may not originate from RGPs but from other types of *Emx1*-CRE labeled progenitors, such as EMX1^+^ intermediate progenitors (IPs) (47, 48). Furthermore, from E12.5 to E13.5, the proportion of ET-PN clones remained low, also indicating the lack of RGP-mediated expansion. Thus, our data does not support the existence of ET-PN fate-restricted RGPs, except if it would be a low proliferative one.

The 12% of E12.5 pure IT-PN clones were found across all types of divisions — proliferative, asymmetric, and terminal divisions—their frequency was very low in proliferative clones (≈ 3%) but constituted ≈ 22% of the neurogenic divisions. At E13.5, their frequency increased significantly in all categories. These results indicate a lineage progression through the early specification of RGPs to IT-PN fates (Figure **5D**). Statistical analysis confirmed that such prevalence of IT-PN clones could not be explained by the death of ET-PNs in Mixed clones (see Methods). Hence, MADM demonstrates the early emergence of IT-PN fate-restricted RGPs.

Next, we investigated the neurogenic decisions dictating the progression of E12.5 multipotent RGPs. For this, we analyzed potential decisions in the daughter cells and the separate subclones (from here after referred as sub-lineages) (Figure **5E-I**). Approximately half of the sub-lineages emerging from the E12.5 RGP population were Mixed sub-lineages. Among the other half, there were almost as many ET-PN sub-lineages as IT-PN sub-lineages. From E12.5 to E13.5, the proportion of IT-PN sub-lineages increased at the expense of decreases in multipotent sub-lineages, and there no changes in ET-PN sub-lineages (Figure **5E**). Hence, ET-PN and IT-PN sub-lineages are specified early (E12.5), and simultaneously and in similar frequencies, but only IT-PN sub-lineages progress through RGPs. In agreement, as with the clones, ET-PN sub-lineages were all small, no larger than 3 cells (Figure **5F**; **S6B**). They were found only among the terminal subclones and within the minority branches of asymmetric subclones (Figures **5H-I**). This again indicated that ET-PNs are not produced from ET-PN fate-restricted RGPs but from the multipotent population, mostly through indirect neurogenesis involving low proliferative progenitors undergoing a maximum of two divisions according to ET-PN subclone size (Figure **5F**).

IT-PN sub-lineages demonstrated progression through RGPs. At E12.5, they showed a wide range of sizes (1-15 PNs) (Figure **5F**; **S6B**), and were found more abundantly in asymmetric and terminal divisions (Figure **5H**). Within the asymmetric divisions, the analysis of the ‘Majority’ branches, which represent the outputs produced by daughter RGPs that emerge with lineage progression, informs of the decisions taken by the parental multipotent RGP population. Half of the ‘Majority’ lineages born from E12.5 asymmetric divisions were Mixed, and the other half were IT-PN lineages (Figure **5H**). This indicated that after any neurogenic division, there is an equal chance for a daughter RGP to maintain multipotency or to become specified for IT-PN production. All in all, the data indicated that rather than priorly existing as an independent population, IT-PN fate-restricted RGPs emerge in half of the neurogenic divisions of multipotent RGPs. By E13.5, IT-PN ‘Majority’ lineages outnumbered Mixed lineages by two-thirds (Figure **5H**). The magnitude of the increase indicates that the daughter RGPs of IT-PN-fated RGPs inherit the restriction, while the daughter multipotent RGPs behave exactly as the parental multipotent RGPs, expanding the pool of IT-PN-fated RGPs in half of their divisions. In this manner, multipotent RGPs, the source of ET-PNs, are depleted with lineage progression (Figure **5K**).

We then examined the laminar distribution of the outputs produced by IT-PN fate-restricted RGPs (i.e.IT-PN subclones ≥4). Notably, nearly all (90%) of IT-PN sub-lineages were translaminar, i.e they produce IT-PNs for both DLs and ULs (Figure **5J**). By E13.5, the fraction of UL IT-PN subclones increased. Thus, IT-PN fate-restricted RGPs display broad layer potential and produce large translaminar outputs through the sequential generation of DL and UL IT-PNs. Overall, the results demonstrate the specification of a subset of IT-PN fate-restricted RGPs, simultaneously to the specification of ET-PN sub-lineages through self-limited intermediate progenitors. The specification as an IT-PN fate restricted RGP correlates with a capacity to generate large translaminar clones. With developmental time, the number of these IT-PN fate-restricted RGPs overgrow, population-wise, the pool of multipotent RGPs, in turn ending the production of ET-PNs (Figure **5K**).

### Small IT-PN and ET-PN lineages from early neurogenic divisions

We next examined the ‘minority’ branches that represent the most immediate neuronal outputs following TM inductions. Notably, at E12.5 most “minority” lineages contained 2-3 cells and thus involved indirect neurogenesis. Moreover, most ‘minority’ lineages were pure (Figure **5H**). This indicated that ET-PN or IT-PN fates are determined, not in the post-mitotic PNs, but at least at the last division in their most immediate progenitor. Hence, rather than direct neurogenesis, the multipotent lineages progress through the initial specification of projection-type restricted progenitors that give rise to small (self-consuming) IT-PN and ET-PN short sub-lineages. Somehow unexpected given the EdU birth dating experiments, we found as many ‘minority’ IT-PN sub-lineages as ET-PN sub-lineages, both at E12.5 (48% ET-PNs vs. 33% IT-PNs) or E13.5 (43% ET-PNs vs. 57% IT-PNs) (Figure **5H**). The numbers were slightly skewed towards ET-PN lineages at E12.5 and IT-PN lineages at E13.5 (Figure **5H**), but could not account for the earlier production of ET-PNs demonstrated with EdU.

To investigate the causes of the sequential production of ET-PNs and IT-PNs demonstrated with EdU, we classify the ‘minority’ outputs in one-cell, two-cell, and three-cell “minority” lineages (Figure **6A**; **S6B**) (11, 49). IT-PN production showed any one-to-three clone size while ET-PN lineages were enriched in two-cell subclones. This suggests distinct modes of proliferation within the sub-lineages. Besides, ET-PN minority subclones were often pairs of the same layer (Figure **6C**). Together, these results strongly suggest that ET-PNs are preferentially generated via IPs derived from multipotent RGPs, which, according to EdU results, undergo terminal divisions at E13.5. In contrasts, over one-third of the IT-PNs produced in the ‘minority’ subclones located in the ULs (Figure **6D**). Since UL-PNs are born at E14.5 onwards, this result indicated that ‘minority’ IT-PN subclones involve progenitors that produce 2-3 PN outputs over a prolonged time window (E12.5-E14.5) and differently from ET-PN IPs. Reported progenitors with such behavior include self-consuming RGPs and other types of intermediate low-proliferative neuronal progenitors (50–52). All in all, the data indicate that small ET-PN and IT-PN subclones involve progenitors that produce outputs with distinctive modes of progression. The precise types of progenitors require further investigation. From the data, we concluded that neurogenic divisions of multipotent RGPs produce pure, small, self-consuming lineages of either ET-PNs or IT-PNs. The divergent progression of the two types of small lineages accounts for the temporal offset between ET-PN and IT-PN birthdates.

**Figure 6.**
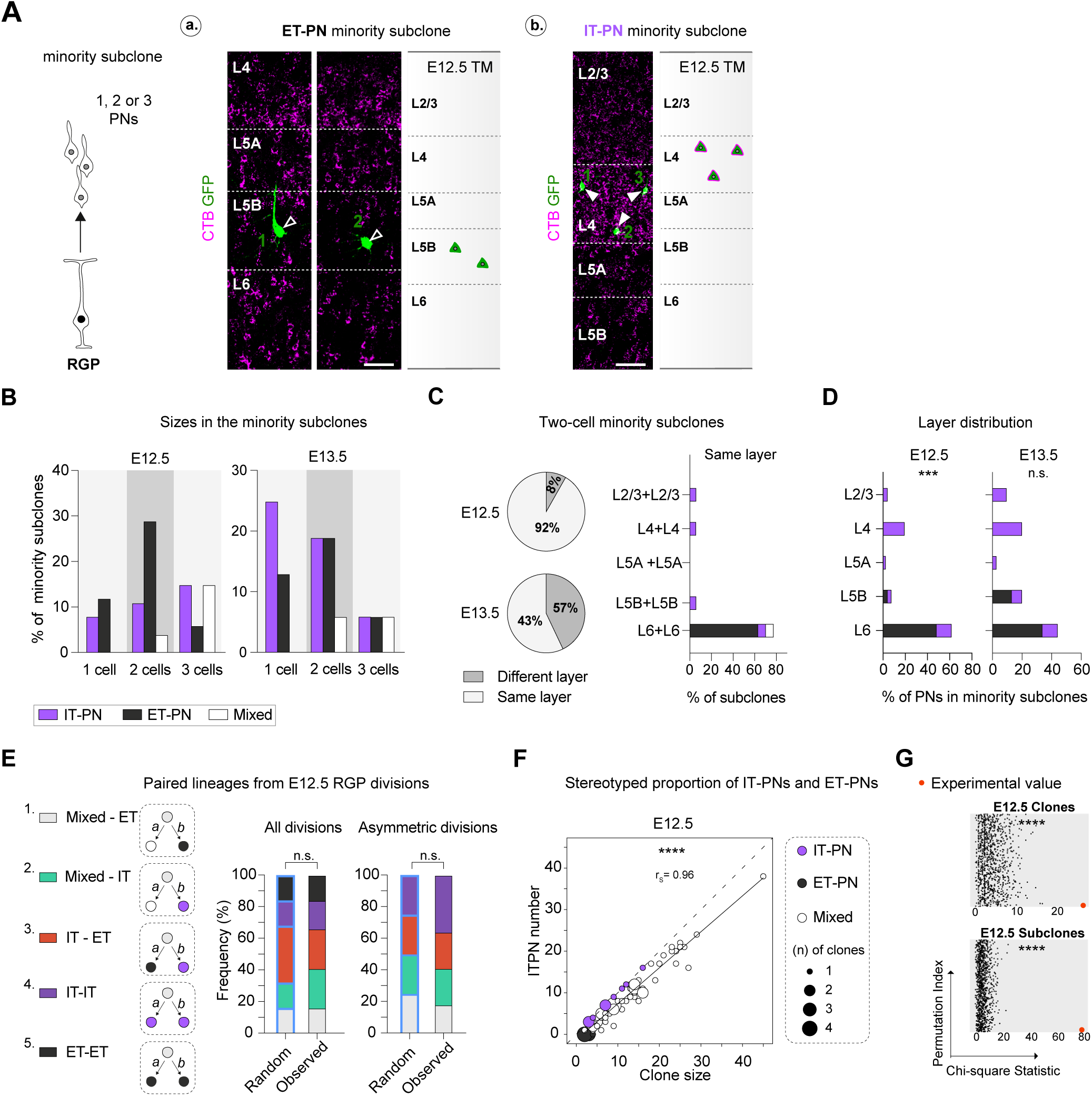
Analysis of IT-PN and ET-PN pure small sub-lineages. (**A**) (Left) Schematic and (Right) Representative images and reconstructions of pure ET-PN (a) or IT-PN (b) minority subclones. CTB^-^PNs are indicated by empty arrowheads, CTB^+^PNs by white arrowheads Scale bars: 50 μm. (**B**) Proportions of pure IT-PN (purple), pure ET-PN (black) and mixed (white) minority subclones containing one, two or three PNs labeled during asymmetric E12.5 or E13.5 MADM divisions. (**C**) (Left) Fraction of two-cell minority subclones with neurons in the same layer. (Right) Layer distribution at P10. (**D**) Differences in the layer distribution between IT-PNs and ET-PNs in the minority subclones. Chi-square test: (E12.5) *** p-value =0.0002, (E13.5) (n.s.) p-value =0.0632. (**E**) (Left) Schematic of the five combinations of paired sub-lineages and (Right) their proportions following TM induction at E12.5 or E13.5. Chi-square test comparing E12.5 and E13.5 distributions to a theoretical random distribution: (All divisions) (n.s.) p-value =0.79, (Asymmetric divisions) (n.s.) p-value =0.75. (**F**) Plot of IT-PN number vs. clone size for clones labeled at E12.5. The bisector (dashed line) indicates IT-PN clones. The black line represents linear regression: y = 0.8646x Clone size – 1.28, where y is IT-PN number. Spearman correlation (r_s_ = Spearman’s rank correlation coefficient): r_s_ (75) = 0.96, **** p-value < 0.0001. (**G**) Association between clone/subclone type (IT-PN, ET-PN, Mixed) and the type of division (P/A/T). A chi-square statistic from the observed data were compared to the chi-square statistics from 1000 random permutations of the dataset. The p-value is calculated as the proportion of permuted chi-square statistics that are greater than or equal to the observed chi-square statistic: **** p-value < 0.0001.

### IT-PN fate-restricted RGPs orchestrate a stereotyped neurogenesis

Our data aligned with previous studies indicating that an initial population of multipotent RGPs gives rise to all PNs in the cortex. However, we found that E12.5 RGPs produce ET-PN and IT-PN sublineages simultaneously not sequentially, and we observed frequencies that challenge the idea of a unique pre-programmed sequence of stereotyped neurogenic divisions.

To evaluate the nature of fate decisions, we compared how the sibling sub-lineages pair with each other. Theoretically, independent decisions in the two daughter cells would produce all possible combinations of paired sub-lineages in random frequencies (Figure **6E**). Conversely, if a multipotent RGP generated ET-PN and IT-PN lineages in a stereotyped manner, IT-PN lineages would never pair with multipotent (Mixed) RGPs. Instead, the predominant combinations should be ET/Mixed and ET/IT pairs. In E12.5 MADM divisions, i.e., the multipotent population, we found all possible combinations of paired sub-lineages in frequencies that support random production, both in the total clones and specifically in the neurogenic asymmetric clones (Figure **6E**) (Methods). Hence, the activation of an IT-PN restriction after the neurogenic division of a multipotent RGP occurs independently of the fate of the small sub-lineages produced in the same division and randomly in the daughter cells. As a result, multipotent RGPs progress through neurogenic sequences that differ at the initial divisions but are stereotyped once an IT-PN fate-restricted RGP emerges. In this scenario, the small and equal frequencies of E12.5 IT-PN and ET-PN clones that we report could be simply the statistical chance result of two sibling cells being independently specified for the same projection-type. For RGPs, there are two choices, multipotency (no restriction) of IT-PN restriction.

The data support a model of corticogenesis in which a random decision, i.e., the activation of an IT-PN restriction in RGPs determines a temporal constrain: the sharp ending of ET-PN production with the extinction of multipotent RGPs. To evaluate whether a random specification renders an overall quantitatively stereotyped process population-wise, we plotted clone size (n) against the number of IT-PNs. This analysis showed a broad range of sizes for Mixed clones (white dots) in both E12.5 and E13.5 populations, with E12.5 clones generally larger. Notably, linear regression analysis revealed a robust relationship between IT-PN content and clone size and demonstrated, on average, a proportional parallel increase in IT-PNs and ET-PNs with clone size (Figure **6F**; **S6C**). This indicated that the production modes we observed render a predictable population, whereby most larger clones are composed of smaller, repeated modular units. Moreover, as in earlier MADM analysis (8), 9 was the most frequent PN unit size of asymmetric clones (Figure **S5E**). Thus, per our linear regression, a canonical E12.5 neurogenic RGP (9 cells) produces an average of 6-7 IT-PNs and 2-3 ET-PNs (Figure **6F**). Finally, probabilistic analysis demonstrated a strong correlation between clone and subclone size and type (ET-PN, IT-PN, or Mixed) (Figure **6G**). This indicated that multipotent, ET-PN, and IT-PN lineages and sublineages each correlate with specific sizes, supporting that proliferation and PN specification are intrinsically linked. Overall, the analysis revealed that, although the neurogenic process does not follow a unique stereotyped initial division, it yields a quantitatively and temporally stereotyped neurogenesis.

### *Pou3f2* drives IT-PN fate

MADM clonal analysis revealed that E12.5 RGPs give rise to IT-PN fate-restricted RGPs. To identify molecular determinants underlying fate restriction specifically in RGPs, we re-analyzed a previously published single-cell RNA sequencing dataset of cortical cells from E12.5 to E15.5 (41). In this study, progenitors were annotated as apical progenitors (APs), containing RGPs, and IPs. POU3 TFs are known to influence UL-PN numbers and identity (53–56), and to regulate proliferation, cell cycle exit, and the balance between direct and indirect neurogenesis in murine cortical progenitors (57). Notably, UMAP embedding of cells clustered by cell type and developmental stage (Figure **7A**; **S7A**), revealed an interesting expression profile for these TFs. *Pou3f1*/*2*/*3* RNA levels were initially absent in early-born PNs, but from E14.5 onwards, they were found in clusters of migrating and immature neurons. In progenitors, most IPs of E14.5-E15.5 cortices, but only a fraction in E12.5-E13.5 cortices, expressed *Pou3f1/2/3*. Most significantly, in APs, *Pou3f2* identified increasingly larger subsets of cells from E12.5 to E15.5, *Pou3f3* showed intermediate levels of expression in all APs invariably across all stages, and *Pou3f1* levels were low at all stages (Figure **7A**; **S7A-B**). Hence, *Pou3f2* expression showed temporal changes expected from an IT-PN restriction in RGPs.

**Figure 7.**
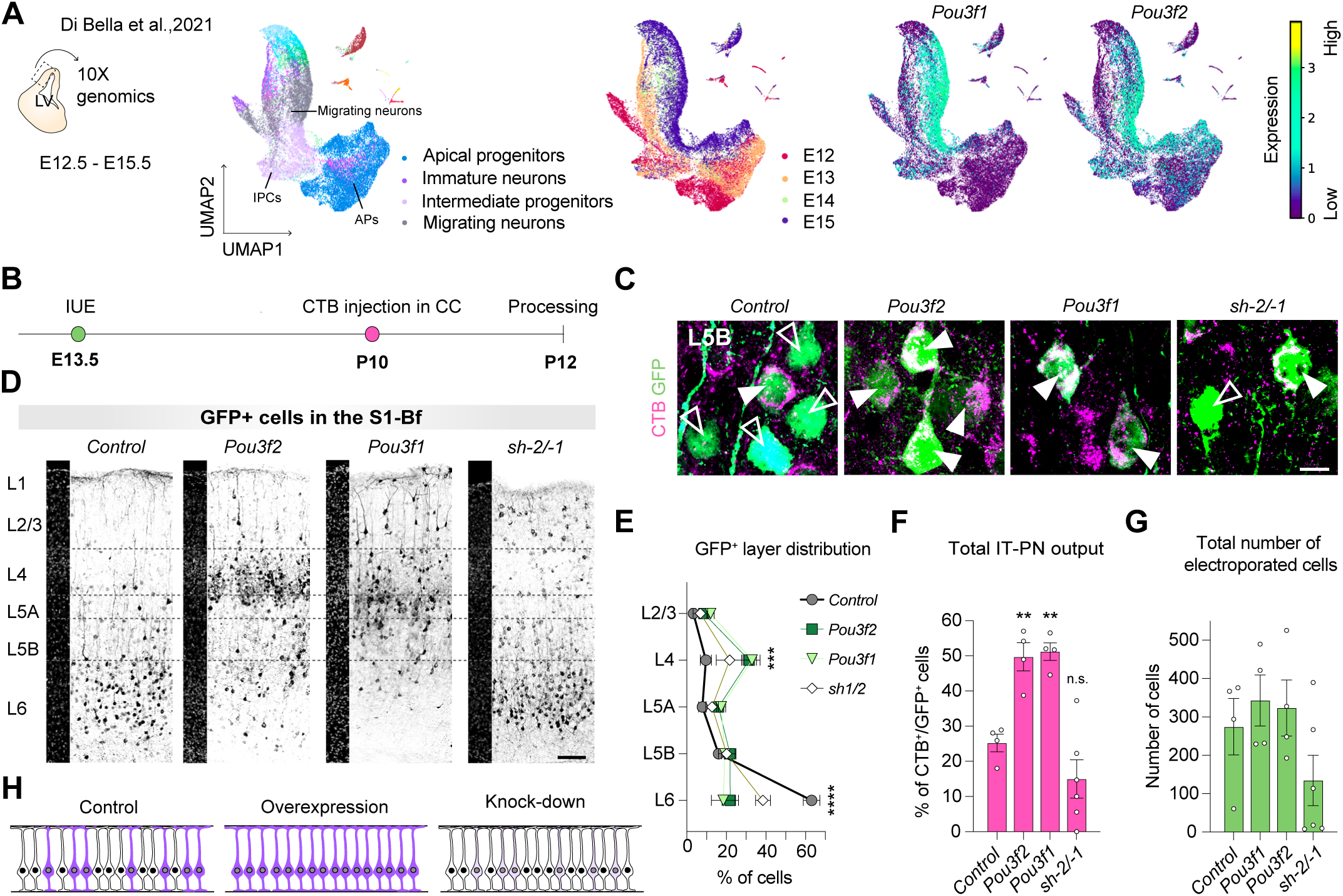
*Pou3f2* regulate IT-PN outputs. (**A**) (Lef) UMAP plots of annotated cell populations across stages (E12.5-E15.5). (Right) Pou3f1/2 expression. (**B**) Experimental design: Control (*Gfp*-), Pou3f2 (*Pou3f2*- plus *Gfp*-), Pou3f1 (*Pou3f1Gfp*-) or shPou3f2/-1 (*shPou3f2*- plus *shPou3f1-* plus *Gfp*-) plasmids were electroporated at E13.5. IT-PNs were identified through CTB injection into the CC at P10, and tissue was collected at P12. (**C**) Electroporated neurons (GFP, green) in L5B positive (IT-PN, white arrow) or negative (empty arrow) for CTB labeling (magenta). Scale bar: 10 μm. (**D**) GFP signal (black) in the S1-Bf following IUE at E13.5 with the corresponding plasmids. Scale bars: 100 μm. (**E**) Layer distribution of electroporated PNs. Two-way ANOVA with Dunnet’s posthoc test: *** p-value _L4 Control vs. L4 Pou3f2_ =0.001, *** p-value _L4 Control vs. L4 Pou3f1_ =0.004, **** p-value _L6 Control vs. L6 Pou3f1 or Pou3f2 or sh2/-1_ <0.0001. (**F**) Percentage of IT-PNs (CTB^+^) over the electroporated population (GFP^+^). Mean ± SEM *(n* = CTB^+^GFP^+^ cells/S1-Bf*, n* = 2 sections/animal; *n* ≥ 3 animals/condition). One-way ANOVA with Dunnett’s posthoc: ** p-value _Control vs. Pou3f2_ = 0.0063, ** p-value _Control vs. Pou3f1_ = 0.0041, p-value _Control vs. sh2/-1_ =0.2489. (**G**) Total GFP^+^ cells in the S1-Bf. One-way ANOVA showed no significant differences. (**H**) Schematic of the VZ in each experimental condition. IT-PN fate restricted RGPs are depicted in purple, multipotent RGPs in white.

To investigate whether POU programs regulate IT-PN production, we performed IUE to overexpress and knockdown *Pou3f2* in E13.5 VZ progenitors (Figure **7B-H**). In these experiments, we also targeted *Pou3f1* to ensure a broad efficient modulation of POU programs, as it is the POU TF with the highest DNA affinity (53). The plasmids included *Gfp* sequences for visualization of electroporated neurons. To identify electroporated IT-PNs, we injected CTB into the CC al P10 and quantified the proportion of CTB^+^GFP^+^ cells at P12 (Figure **7B**).

In the control condition, most of the E13.5 GFP^+^ outputs were comprised of CTB^-^ PNs (ET-PNs) that migrate to the DLs (Figure **7C-F**). Overexpression of *Pou3f2* or *Pou3f1* increased the proportion of CTB^+^GFP^+^ cells (IT-PNs) from 20% to approximately 50% of the GFP^+^ population (Figure **7F**) and shifted the layering of electroporated cells towards the ULs (Figures **7E**). This data demonstrated that POU programs promote IT-PN specification. Conversely, *Pou3f2*/-*1* knockdown reduced the overall PN output (Figure **7G**), in agreement with *Pou3f* regulation of the proliferation of VZ progenitors (57), it broadened GFP^+^ PN distribution across layers (Figure **7E**), and equalized the production of CTIP2^+^- ET-PNs and DL-IT-PNs (CTB^+^), decreasing it from 3:1 to 1:1 ratio (Figure **S7C-D**). These alterations supported that in the absence of *Pou3f*, multipotent RGPs produce only small lineages (Figure **7H** and **8**) and not the IT-PN fate-restricted RGPs that would continue to proliferate and dilute the GFP plasmids. All in all, the experiments indicated the necessity of *Pou3f2* for the specification of IT-PN fate-restricted RGPs and suggested that IT-PN fate is coupled with the increased proliferative capacity of RGPs.

**Figure 8.**
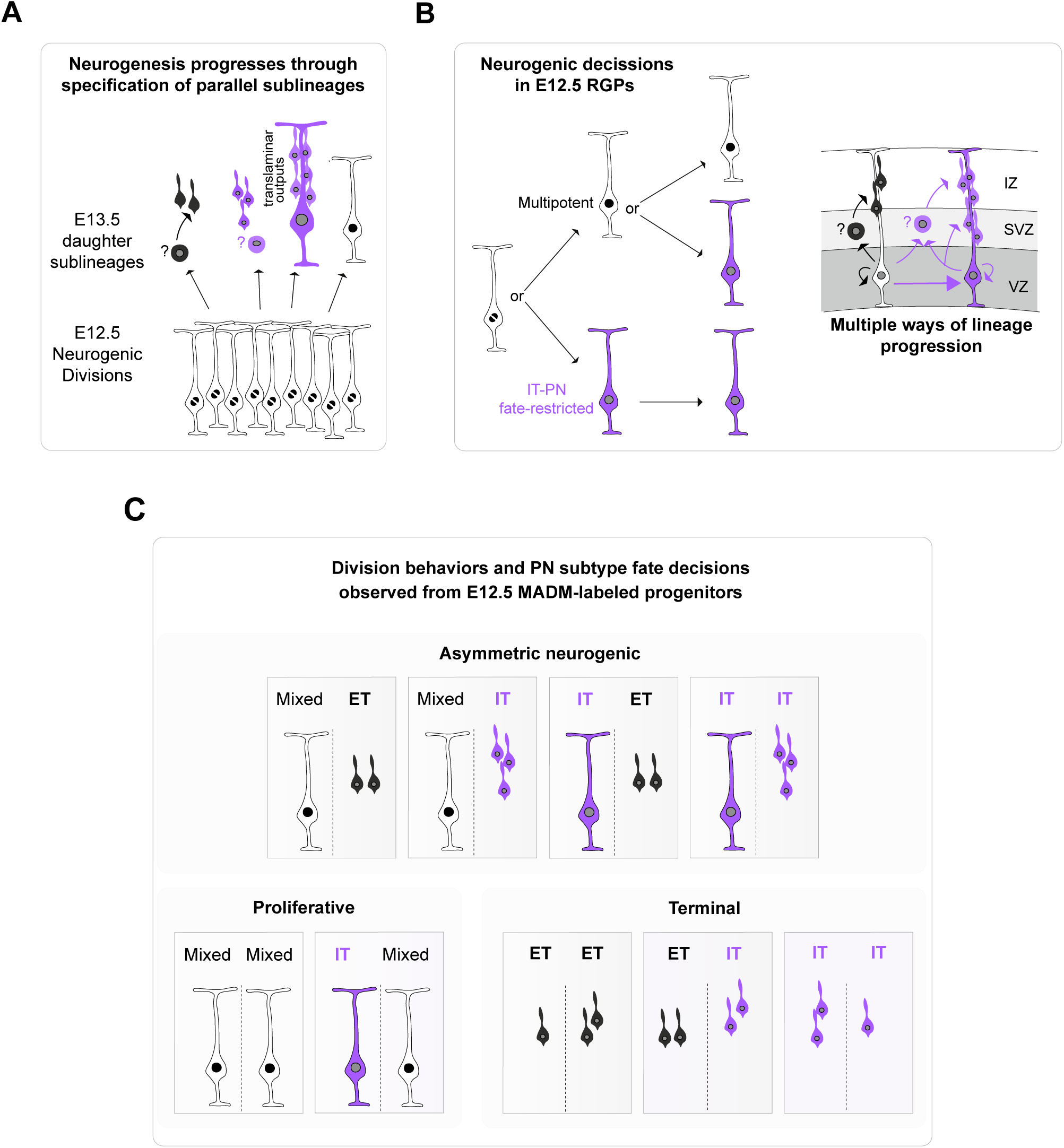
Model illustrating RGP lineage progression. IT-PN sub-lineages are depicted in purple, ET-PN sub-lineages in black, and multipotent RGPs in white. **(A)** Neurogenesis progresses through fate-restricted sub-lineages. Upon entering neurogenesis at E12.5, four types of subclones emerge from the multipotent RGP population: small self-consuming ET-PN (moist frequent size 2), small IT-PN lineages (most frequent size 3), large translaminar RGP mediated IT-PN subclones, and multipotent (ET-PN and IT-PN) subclones. **(B)** In the canonical assymetric divisions generating a Majority and a minority subclones, we observe that neurogenic multipotent RGPs divide and generate RGP daughters that with equal probability are i) IT-PN fate-restricted or ii) maintain multipotency. This decision is independent of the fate of the minority lineage. **(C)** Scheme illustrating the types of divisions and sub-lineage combinations traced in this study with MADM clonal analysis.

## DISCUSSION

We herein demonstrate that mouse RGPs are not a uniform population but consist of at least two main types: multipotent RGPs and IT-PN fate-restricted RGPs, the product of lineage progression. Importantly, genetic MADM labeling at E13.5 demonstrates that IT-PN fate-restricted RGPs emerge from E12.5 divisions and progress to generate translaminar lineages with differentiation. We also reveal that E12.5 multipotent RGPs appear not pre-programmed at the individual level. Upon entering the neurogenic phase, the asymmetric division of an E12.5 RGP produces with no specific preference and in similar frequencies (i) multipotent or IT-PN fate-restricted RGPs and (ii) small pure ET-PN or IT-PN lineages (1-3 PNs) (Figure **8A** and **8B**). Nevertheless, the whole process of corticogenesis is stereotyped. Intrinsic to this early freedom and an apparent lack of order, the emergence of IT-PN fate-restricted RGPs introduces a single constraint that yields a temporarily and quantitatively stereotyped ordered PN production.

The existence of IT-PN fate-restricted radial glial progenitors (RGPs) is a key finding of this study. In the cortex, non-clonal fate mapping experiments identified RGP subsets expressing different transcription factors (e.g., Lhx2 and Fezf2), with Fezf2^+^ RGPs specifically generating ET-PNs (17). Reconciling these observations with our data suggests that *Fez2+* might identify multipotent RGPs while the absence of *Fezf2* expression might define IT-PN fate-restricted RGPs. Likewise, our data on *Pou3f2* expression and function indicates that this TF molecularly defines IT-PN fate-restricted RGPs, supporting that the diversity of PN types is already reflected in the heterogeneity of RGPs at the transcriptomic level, at least to a certain level. In the context of neurodevelopmental diseases, *POU3F2* variants are associated with bipolar disorders and schizophrenia, and in mice, loss of function *Pou3f2* mutations produce microcephaly (57, 58). Both phenotypes could be related to failures in the specification and production of IT-PN lineages. Indeed, IT-PNs are the most expanded and diversified PN type across mammals and are highly affected in neurodevelopmental diseases (20, 59, 60). Conversely, it is worth speculating that multipotent RGPs can only produce self-consuming sublineages and that the specification of IT-PN fate-restricted RGPs may have enabled the expansion of PN production and of the mammalian brain over evolution.

Another important aspect of our data is the mode of lineage progression. Our data supports the prevailing view that all mouse cortical PNs originate from an initially homogeneous population of multipotent RGPs (18–21, 41, 61–68) but challenge the notion that cortical PNs emerge from a pre-determined sequence of changes in RGP competence. The early stochasticity we observe in the first neurogenic decisions provides evidence against an invertebrate-like pre-defined program of precise neurogenic instructions governing mammalian corticogenesis (69–71). Pioneer MADM studies already indicated a certain heterogeneity in RGP outputs. Gao et al., and subsequently Llorca et al, found that not all MADM clones contained PNs of all layers (L2-L6) but exhibited variable layer configurations. We now confirm that multipotent RGPs progress in their lineage through multiple sequences that collectively account for a stereotyped PN production.

We show that IT-PNs and ET-PNs birth only partially overlap and that many late-born (E14.5) IT-PNs migrate to the DLs, which was also observed in the pioneer birth-dating studies in the mouse cortex (45). Conversely, more recent studies showed that isochronally born neurons adopt diverse layer identities (72). Our data demonstrates that, indeed, the birth time alone does not determine the PN layer position. We show that the hierarchical superimposition of the projection-subtype decision over a broad layer-identity potential is transformed in a specialized RGP sublineage of protracted progression when compared to multipotent RGPs.

We demonstrate that proliferation and specification are intrinsically linked, as evidenced by the significant correlation between the size and projection subtype of the lineages (Figure **6G**). Our findings unveil that multipotent RGPs are not committed but PN specification of their outputs occurs in the progenitors they generate. ET-PN and IT-PN specification is achieved through distinct progenitor types that emerge and are specified simultaneously. For ET-PNs, progenitors seem to respond to the classical IPs. For IT-PNs, we might be facing two or more types of progenitors enabling the production of translaminar two- to-three PN outputs through asymmetric divisions. On one hand RGPs, on the other, another type o low proliferative progenitors in the small sub-lineages. By all means, the two types could be related. In fact, slow-cycling RGPs have previously been associated with the generation of ULs (73), making it plausible that all IT-PNs are produced from an RGP, whether low proliferative RGPs or IT-PN-restricted canonical RGPs. Other progenitor types could account for IT-PN small lineages: (i) RGP-derived apical intermediate progenitors, also called short neural precursors (SNPs), which display a slow cell cycle kinetic and predominantly produce L4 PNs (50–52); (ii) outer SVZ RGPs (oRGs), also referred to as basal RGs (bRGs) (74–76), which have been closely associated with IT-PN production in humans and non-human primates (77, 78); (iii) IPs cycling more than once, which are nonetheless sparse in mice (49, 79). Regardless, we show data indicating that the sequentially of ET-PN and IT-PN birth is rooted in the distinct progression of their respective sublineages. Indeed, Notably, POU3F2 and POU3F1 TFs are known to regulate cell proliferation and cell-cycle exit of mouse RGPs (53, 55, 57).

Finally, concerning small lineages, it is worth mentioning the possible implications of the observation of terminal clones and the inclusion of these data in the analysis. First and foremost, observations of terminal clones should be interpreted with caution. Using the MADM protocol, we genetically label Emx1-expressing progenitors. While in asymmetric and proliferative clones, size indicates RGPs, terminal clones represent an undefined category of self-consuming Emx1^+^ progenitors that could include low proliferative RGPs and some of the above-mentioned progenitors of the small lineages (48). Noteworthy, the lineages we observe in the terminal clones are indistinguishable from those in the minority branches. Hence, the terminal divisions could represent two independently fated small lineages generated from a neurogenic division of multipotent RGPs. Regardless of the specific scenario, our finding supports the frequent production of small lineages through low-proliferative progenitors during early neurogenesis, an issue that requires further investigation beyond the scope of the present study. All in all, our results highlight the biological relevance of progenitor diversity in lineage progression and the capacity of these progenitors to carry projection-subtype PN identity, a concept suggested in recent studies (17, 41, 80).

In summary, we discovered an RGP lineage progression and early modes of neurogenic divisions that challenge the classical view of a canonical multipotent RGP type progressing through a unique neurogenic sequence. Instead, our data support a model in which the early emergence of parallel early fated sub-lineages and most specifically, we reveal that the establishment of an IT-PN fate-restriction in RGPs on top of a broad layer-identity potential, allows the expansion of the multipotent lineage and generates a globally stereotyped process. The findings have broad implications for neurodevelopmental disorders and evolution.

## Acknowledgments

We thank M. Caouyette for the plasmid construction for *Pou3f1* overexpression; C. Varela-Martínez for help with the code for graphical analysis; all members from the Nieto’s lab for comment on the manuscript, specially to F. Martín for the insightful discussions; J.C. Oliveros and J.A. García from the computational service of the CNB for help with the analysis of RNAseq dataset, C.O. Sorzano for the help with statistical analysis, and the service of Advance Optical Microscopy of the CNB for their technical advice.

## Funding

I.V.M holds a fellowship funded by MCICIU (PRE-2018-083376), the work was funded by PID2020-112831GB-I00 funded by MCIN/AEI /10.13039/501100011033.

## Author contributions

Conceptualization: IVM, SH, MN

Methodology: IVM, AVR, JGM, SH, MN

Investigation: IVM, AVR, MN

Visualization: IVM, MN

Supervision: SH, MN

Writing – original draft: IVM, MN

Writing – review & editing: IVM, JGM, AVR, SH, MN

## Competing interests

Authors declare that they have no competing interests.

## Data and materials availability

All data are available in the main text or the supplementary materials.

## Notes

https://doi.org/10.5281/zenodo.14609057

### Competing Interest Statement

The authors have declared no competing interest.

