## Supplemental material for "Early Emergence of Projection-subtype fate-restricted Radial Glial Progenitors Orchestrates Neocortical Neurogenesis"

#### SUPPLEMENTARY FIGURE LEGENDS

**Figure S1. CTB injections to investigate the developmental dynamics of callosal projections in the S1-Bf, related to Figure 1**

(**A** and **B**) NeuN (green), CTB-555 (magenta) and DAPI (blue) signals in the ULs of brains injected and analyzed at the indicated times. Refer to main figures 1E and 1F.

(**C**) CTB labeled CPNs (magenta) following P0, P5, P16, or P30 CC injections (left to right, respectively).

(**D-G**) Percentages of CPNs quantified as CTB-555$^+$NeuN$^+$ cells over 50 NeuN$^+$ per layer, following CTB injections in the CC at P0 (**D**), P5 (**E**), P16 (**F**), or P30 (**G**) Mean ± SEM ($n$ = 500 cells, $n$ = 2 sections/animal; $n \geq 4$ animals). One-way ANOVA showed significant differences in CTB$^+$NeuN$^+$ percentages across layers at each evaluated point (**** p-value <0.0001). Corresponding posthoc with Tukey's test below each panel.

**Scale Bars:** 50 μm (A, B and C)

**Figure S2. The striatum is a preferential target for a subset of early PNs with developmental callosal axons, related to Figure 1**

(**A**) Schematic illustrating the analysis of PNs projecting dually to both the CC and to the striatum (Str): (a) CTB-555 (magenta) was injected into the CC at P3 followed by CTB-488 (green) injections into the Str at P5; (b) representation of a PN projecting ipsilaterally to the Str (iStr) and callosally; (c) representation of a CPN projecting to the contralateral Str (cStr).

(**B**) CTB-488 signal confined within the dorsal-Str two h post-injection.

(**C**) CTB-488 labeling of S1-Bf PNs projecting to the ipsilateral (iStr-PN, left) or contralateral (cStr-PN, right) Str following injections as outlined in (A).

(**D**) Percentage of CTB-488$^+$ iStr-PNs and cStr-PNs following injections into the Str as in (A). Mean ± SEM ($n$ = CTB-488$^+$ cells in the S1-Bf, $n$ = 2 sections/animal, $n$ = 3 animals).

(**E** and **F**) S1-Bf cortices ipsilateral (E) or contralateral (F) to the striatal injections after injections as described in (A). High-magnifications in (E) show L5A and L5B dually labeled cells (white arrowheads) and single-labeled cells (empty arrowheads).

(**G**) Percentages of CTB-488$^+$ (striatal injection) over the CTB-555$^+$ (callosal injection) cells per layer. Mean ± SEM ($n$ = 500 cells/hemisphere, $n$ = 2 sections/animal, $n$ = 3 animals).

**Scale Bars:** 500 μm (B), 50 μm (C), 150 μm (high magnification, E), 300 μm (low magnification, E and F).

**Figure S3. Birthdate analysis of IT-PNs using EdU injections, related to Figure 3**

(**A**) Total EdU$^+$ cells in the S1-Bf at P10 following intraperitoneal EdU injections at the indicated stages. Mean ± SEM ($n$ = EdU$^+$/S1-Bf, $n$ = 2 sections/animal, $n$ = 3 animals/stage). One-way ANOVA test with Tukey's posthoc (comparisons were made between consecutive stages): ** p-value $_{E12.5 \text{ vs } E13.5}$ =0.006.

(**B**) Proportion of cells born in the S1-Bf at the indicated stages per layer. Data were calculated using the averages of the total number of newborn cells per embryonic stage: ($n$= EdU$^+$ per S1-Bf, $n$= 2 sections/animal, $n$=3 animals/stage).

(**C**) Experimental design (top) and high-magnification image (bottom) showing EdU (green), CTB (magenta) and NeuN (white) co-labeling.

(**D**) Proportion of neuronal (NeuN$^+$) and non-neuronal (NeuN$^-$) newborn cells per layer labeled with EdU at E14.5. Mean ± SEM ($n$ = EdU$^+$/S1-Bf , $n$ = 2 sections/animal; $n$ = 3 animals).

(**E**) Percentage of IT-PNs (CTB$^+$) calculated over EdU$^+$ labeling (bars in magenta) or EdU$^+$NeuN$^+$ labeling (bars in gray). EdU injections were performed at E14.5 and CTB injections into the CC at P5. Mean ± SEM ($n$ = CTB$^+$EdU$^+$ or CTB$^+$EdU$^+$ NeuN$^+$/S1-Bf, $n$ = 2 sections/animal; $n$ = 3 animals). Two-way ANOVA with Šidak's posthoc test (CTB$^+$EdU$^+$ vs. CTB$^+$EdU$^+$NeuN$^+$): * p-value $_{L2/3}$ =0.427, * p-value $_{L4}$ =0.0154, ** p-value $_{L5A}$ =0.0023, ** p-value $_{L5B}$ =0.0022, (n.s.) p-value $_{L6}$ =0.9894.

(**F**) EdU (green) and CTB-555 (magenta) labeling in a ROI within the S1-Bf following EdU intraperitoneal injections at the indicated stages. Cortical layers are outlined with white dashed lines.

(**G**) CTB-555 (magenta) and DAPI (blue) signals in the S1-Bf ULs following EdU injections at E11.5, E12.5 and E13.5. Absence of EdU signal (green) indicates no UL-PN generation during these stages.

**Scale bars:** 20 μm (C, G), 100 μm (F).

**Figure S4. IUE confirmed that ET-PNs are only generated during early neurogenesis, related to Figures 3 and 4**

(**A**) Schematic of the experimental design. Dividing progenitors in the VZ were electroporated at E13.5, E14.5, or E15.5 with a *Gfp*-encoding plasmid, followed by analysis at P12.

(**B**) Layer distribution of $GFP^+$ cells in the S1-Bf at P12. Mean ± SEM ($n$ = $GFP^+$ cells/S1-Bf, $n$ = 2 sections/ animal; $n$ = 3 animals/embryonic stage).

(**C**) GFP signal following IUE at E14.5. Green boxes indicate regions of interest: (1) ipsilateral IC and Th, (2) ipsilateral dorsal-Str and WM, (3) contralateral dorsal-Str and WM. Corresponding high magnification images are shown in panels (E) for E13.5, (F) for E14.5, and (I) for E15.5.

(**D**) (Left) High-magnification image and (Right) 3D Imaris reconstruction of an L5A PN extending a dual projection to the CC and the ipsilateral Str.

(**E**, **G,** and **I)** S1-Bf of P12 brains following IUE at E13.5 (E), E14.5 (F), or E15.5 (G),. Cortical layers are outlined by dashed lines.

(**F**, **H** and **J**) High-magnification images of the regions indicated in (C) following IUE at E13.5 (F), E14.5 (H), and E15.5 (J).

**Scale bars:** 1 mm (C), 150 μm (D), 100 μm (E, G, and I), 250 μm (F, H and J)

Abbreviations: PN: Projection neuron, IP: Intermediate Progenitor, RGP: Radial Glial Cell, LV: Lateral Ventricle, VZ: Ventricular Zone, SVZ: Subventricular Zone, CC: Corpus Callosum, IC: Internal Capsule, Th: Thalamus, Str: Striatum, WM: White Matter.

**Figure S5. MADM-based clonal analysis, related to Figure 5**

(**A**) Schematic of MADM labeling. Males and females of *Emx1-CreERT2*$^{+/-}$ and *MADM11*$^{GT/TG}$ genotypes were used indistinctly.

(**B**) Consecutive brain sections of Proliferative and Terminal G2-X clones. Each clone contains a magenta (tdTomato) and a green (GFP) subclone. PN numbers/subclone in corresponding colors. Right-most panels: schematics of the reconstructed clones. Scale bar: 100 μm.

(**C**) Proportion of clones in each category following TM induction at E12.5 or E13.5. Chi-square test: p-value=0.0915. Bold numbers represent the number of clones per category.

(**D-F**) Clone sizes for Proliferative (D), Asymmetric (E), and Terminal (F) clones (box, median with 25-75 percentiles; whiskers, minimum and maximum values). Dots represent neurons per individual clone (n = number of clones per category and stage indicated in C). Mann-Whitney test (E12.5 vs. E13.5): (D) p-value =0.0655, (E) * p-value =0.037, (F) p-value =0.8672.

(**G**) Percentage of single-color/orphan clones. The probability of death is inferred as the proportion of single-color (tdTomato$^+$ or GFP$^+$) clones in the population (15.5%).

(**H**) DL to UL ratio in asymmetric clones. Data represent the median (E12.5 n=27 clones, E13.5 n=16 clones). Unpaired t-test showed non-significant differences.

**Figure S6. MADM analysis of IT-PN, ET-PN and Mixed clones and subclones, related to Figures 5 and 6**

(**A**) Proportion of CTB$^+$ and CTB$^-$ neurons per layer in MADM$^+$ and NeuN$^+$ populations. Chi-square test: (L2/3) **** p-value $_{\text{E12.5 MADM}^+ \text{ vs P5 NeuN}^+}$ <0.0001, *** p-value $_{\text{E13.5 MADM}^+ \text{ vs P5 NeuN}^+}$ =0.0003. Proportions of CTB$^+$NeuN$^+$ were calculated using the data in Figure 1.

(**B**) Size distribution of subclones with two or more PNs.

(**C**) Plot of IT-PN number vs. clone size for clones labeled at E13.5. The bisector (dashed line) indicates IT-PN clones. The black line represents linear regression: y = 0.8955x Clone size – 0.7, where y is IT-PN number. Spearman correlation ($r_s$ =Spearman's rank correlation coefficient): $r_s$ (33) = 0.93, **** p-value <0.0001.

**(D)** Cumulative frequency of asymmetric clones with increasing IT-PN Bin-percentages (Bin=10). Intersection with dashed lines represents the median (E12.5 *n* = 27 clones, E13.5 *n* = 16 clones).

**Figure S7. Analysis of the expression and functions of POU factors in IT-PN production, related to Figure 7**

**(A)** (Left) UMAP plots of annotated cell populations across stages (E12.5-E15.5). (Right) *Pou3f3* expression. Dataset obtained from Di Bella et al. 2021.

**(B)** Immunostaining of E13.5 and E15.5 coronal brain slices for POU3F1 (green) and EOMES (magenta). High-magnification areas indicated with white boxes.

(**C**) (Left) Representative images of L5 PNs after E13.5 IUE with *CAG-GFP* (control) or *shPou3f1* and *shPou3f2* (knockdown) plasmids. GFP is shown in green, CTB in magenta and immunolabeling against CTIP2 in cyan. (Right) Ratio of $Ctip2^{+}$ET-PNs to DL-IT-PNs in control and knockdown electroporated cells (*n* = $Ctip2^{+}$ET-PNs / DL-IT-PNs number, *n* = 2 sections/animal, *n* = 3 animals).

**(D)** (Left) Experimental design for shRNA validation in N2A cells: indicated plasmids were transfected and RNA levels quantified via qRT-PCR. A Scramble shRNA was used as control condition. Five and four candidates were tested against Pou3f1 and Pou3f2, respectively. (Right) Quantification of mRNA expression in the indicated conditions. Mean ± SEM (n = 5 independent cultures). One-way ANOVA with Dunnnett's posthoc: (Pou3f1) ** $p\text{-value}_{\text{Scramble vs Sh1}}$ =0.0078, **** $p\text{-value}_{\text{Scramble vs Sh2}}$ < 0.0001, * $p\text{-value}_{\text{Scramble vs Sh3}}$ =0.0154, ** $p\text{-value}_{\text{Scramble vs Sh4}}$ =0.0016, * $p\text{-value}_{\text{Scramble vs Sh5}}$ =0.0488, (Pou3f2) * $p\text{-value}_{\text{Scramble vs Sh1}}$ =0.0121, * $p\text{-value}_{\text{Scramble vs Sh2}}$ =0.0254, ** $p\text{-value}_{\text{Scramble vs Sh4}}$ =0.0033.

**Scale bars:** 300 µm (low magnification in B), 30 µm (high magnification in B), 10 µm (C).

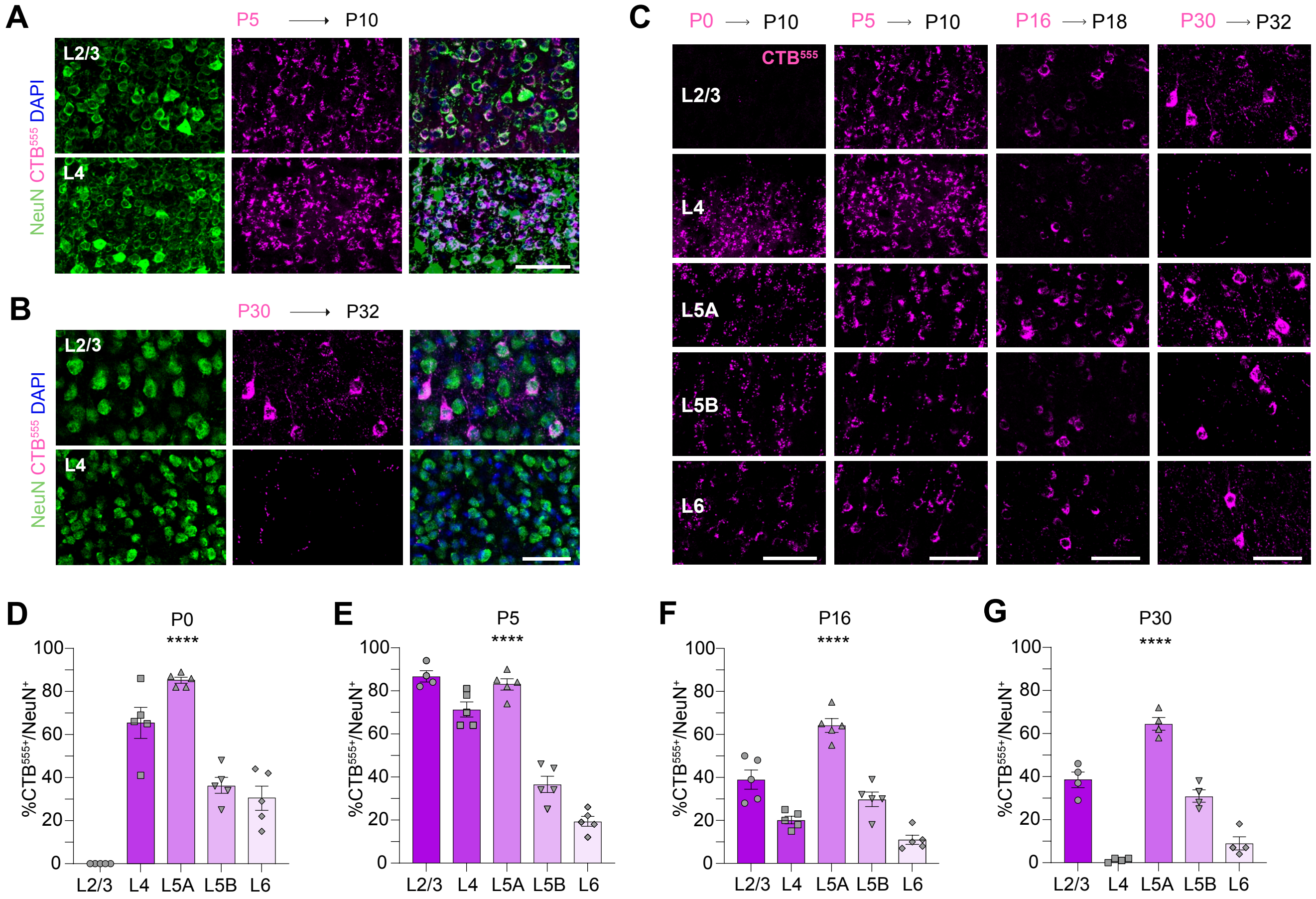


**D**

| | L2/3 | L4 | L5A | L5B | L6 |
|---|---|---|---|---|---|
| L2/3 | *n=5* | | | | |
| L4 | **** <0.0001 | *n=5* | | | |
| L5A | **** <0.0001 | * 0.0362 | *n=5* | | |
| L5B | *** 0.0001 | ** 0.014 | **** <0.0001 | *n=5* | |
| L6 | *** 0.0009 | *** 0.0002 | **** <0.0001 | n.s 0.8722 | *n=5* |

**E**

| | L2/3 | L4 | L5A | L5B | L6 |
|---|---|---|---|---|---|
| L2/3 | *n=5* | | | | |
| L4 | * 0.0225 | *n=5* | | | |
| L5A | n.s 0.9183 | n.s 0.088 | *n=5* | | |
| L5B | **** <0.0001 | **** <0.0001 | **** <0.0001 | *n=5* | |
| L6 | **** <0.0001 | **** <0.0001 | **** <0.0001 | ** 0.0057 | *n=5* |

**F**

| | L2/3 | L4 | L5A | L5B | L6 |
|---|---|---|---|---|---|
| L2/3 | *n=5* | | | | |
| L4 | ** 0.0034 | *n=5* | | | |
| L5A | *** 0.0001 | **** <0.0001 | *n=5* | | |
| L5B | n.s 0.2731 | n.s 0.2368 | **** <0.0001 | *n=5* | |
| L6 | **** <0.0001 | n.s 0.2731 | **** <0.0001 | ** 0.0034 | *n=5* |

**G**

| | L2/3 | L4 | L5A | L5B | L6 |
|---|---|---|---|---|---|
| L2/3 | *n=4* | | | | |
| L4 | **** <0.0001 | *n=4* | | | |
| L5A | **** <0.0001 | **** <0.0001 | *n=4* | | |
| L5B | n.s 0.3734 | **** <0.0001 | **** <0.0001 | *n=4* | |
| L6 | **** <0.0001 | n.s 0.343 | **** <0.0001 | *** 0.0002 | *n=4* |

**Figure S1**_*Varela-Martínez et al.,*

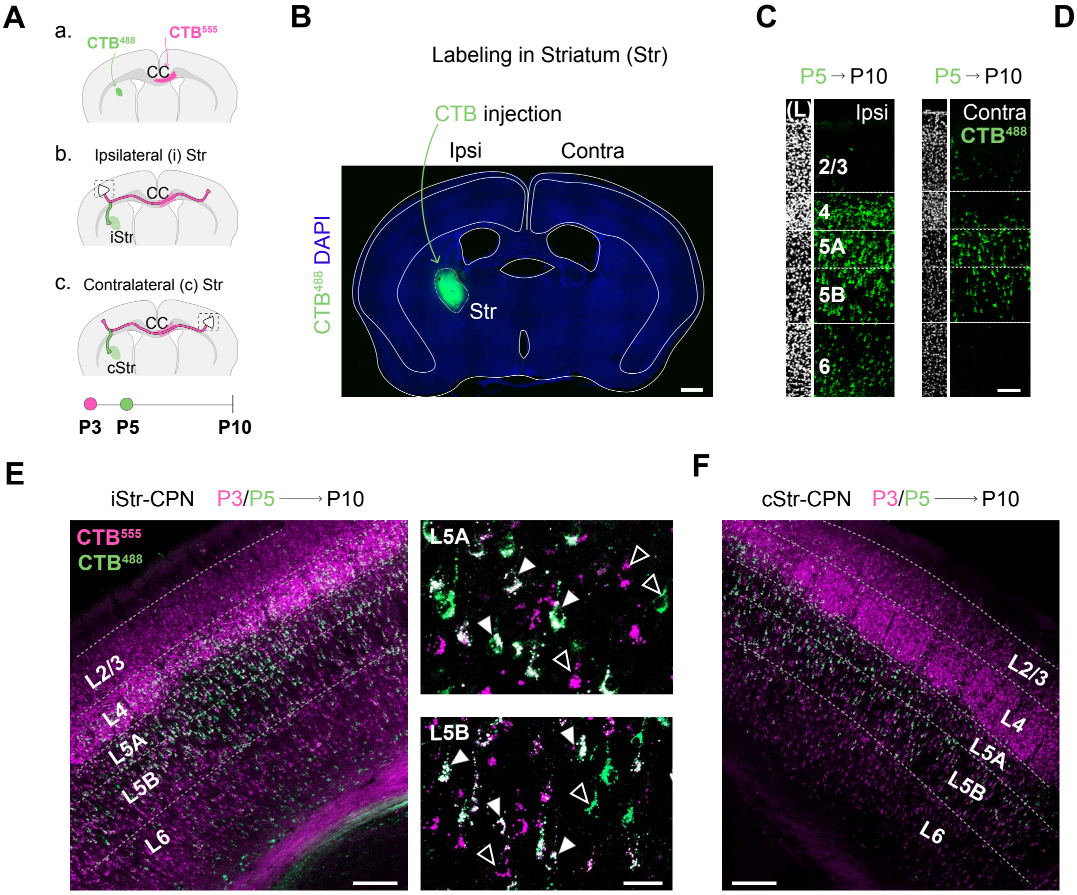


**G**

% of CTB488+/CTB555+ cells

| | iStr-CPN | cStr-CPN |
|---|---|---|
| L2/3 | 5.7% ± 2.3 | 20% ± 14.2 |
| L4 | 18.7% ± 2.2 | 10.7% ± 3.8 |
| L5A | 54% ± 7.6 | 50.7% ± 2.9 |
| L5B | 45.3% ± 8.7 | 33.7% ± 4.9 |
| L6 | 17.3% ± 2.6 | 10% ± 4.6 |

**Figure S2**_*Varela-Martínez et al.,*

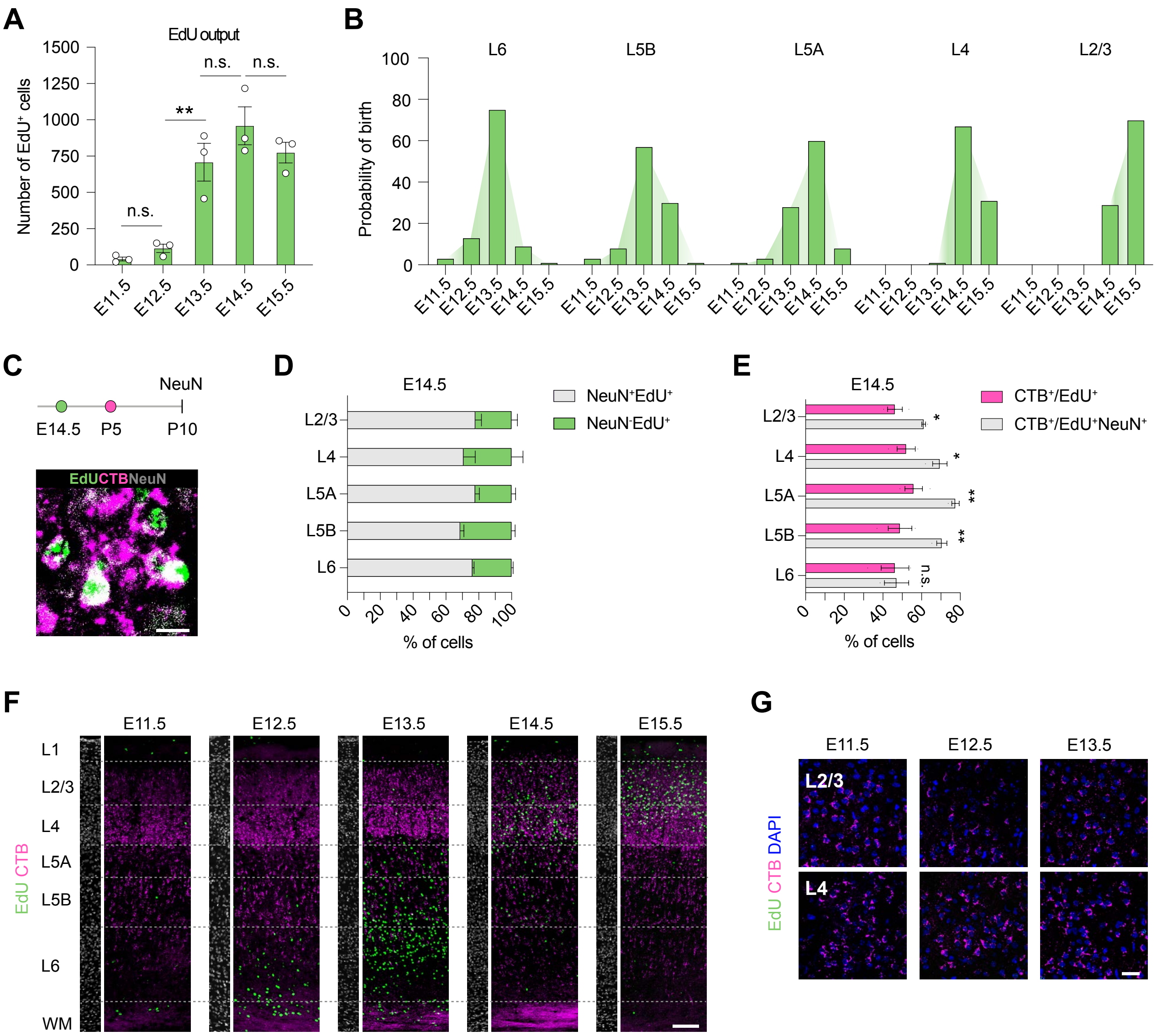


**Figure S3**_*Varela-Martínez et al.,*

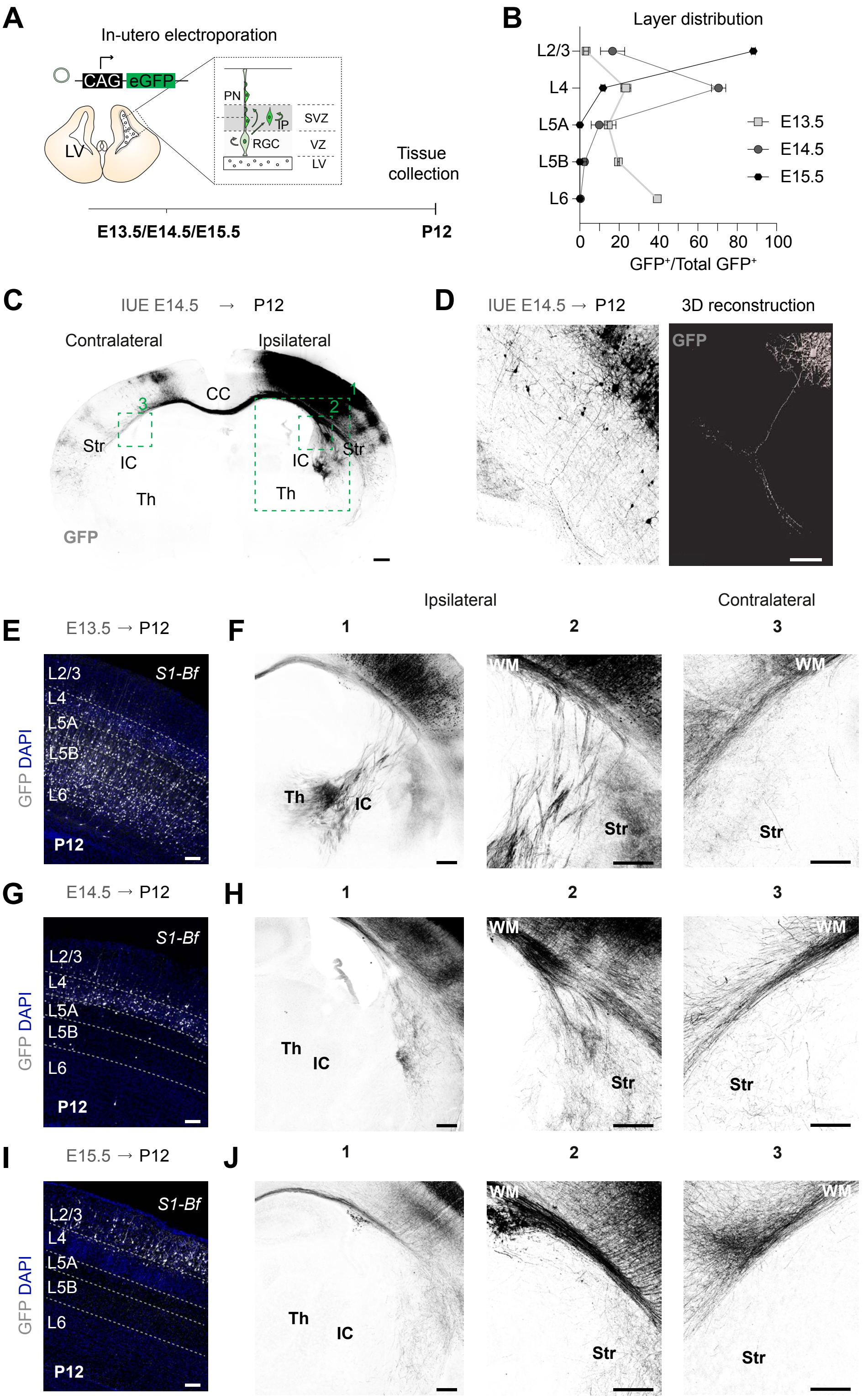


**Figure S4**_*Varela-Martínez et al.,*

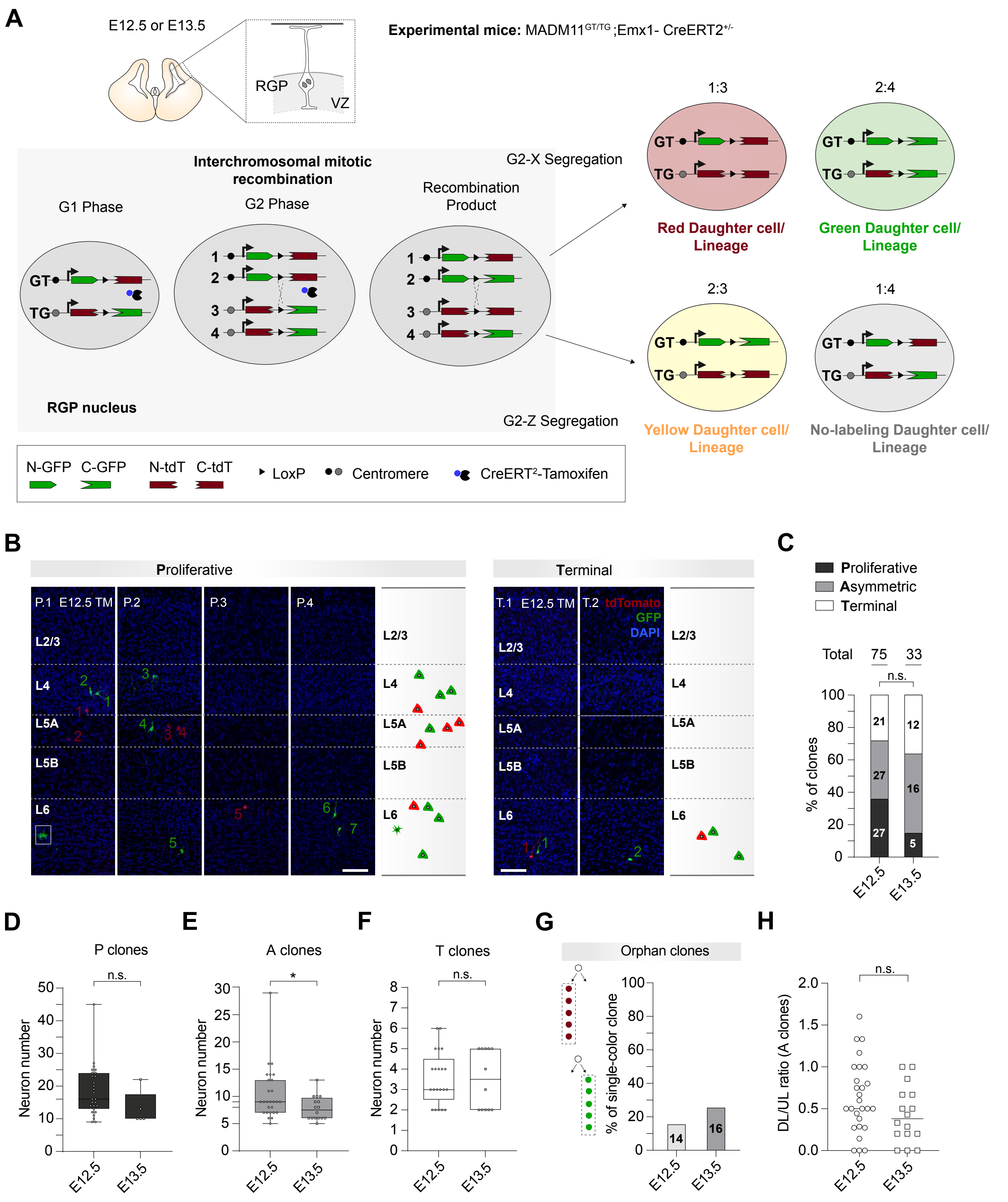


**Figure S5_***Varela-Martínez et al.,*

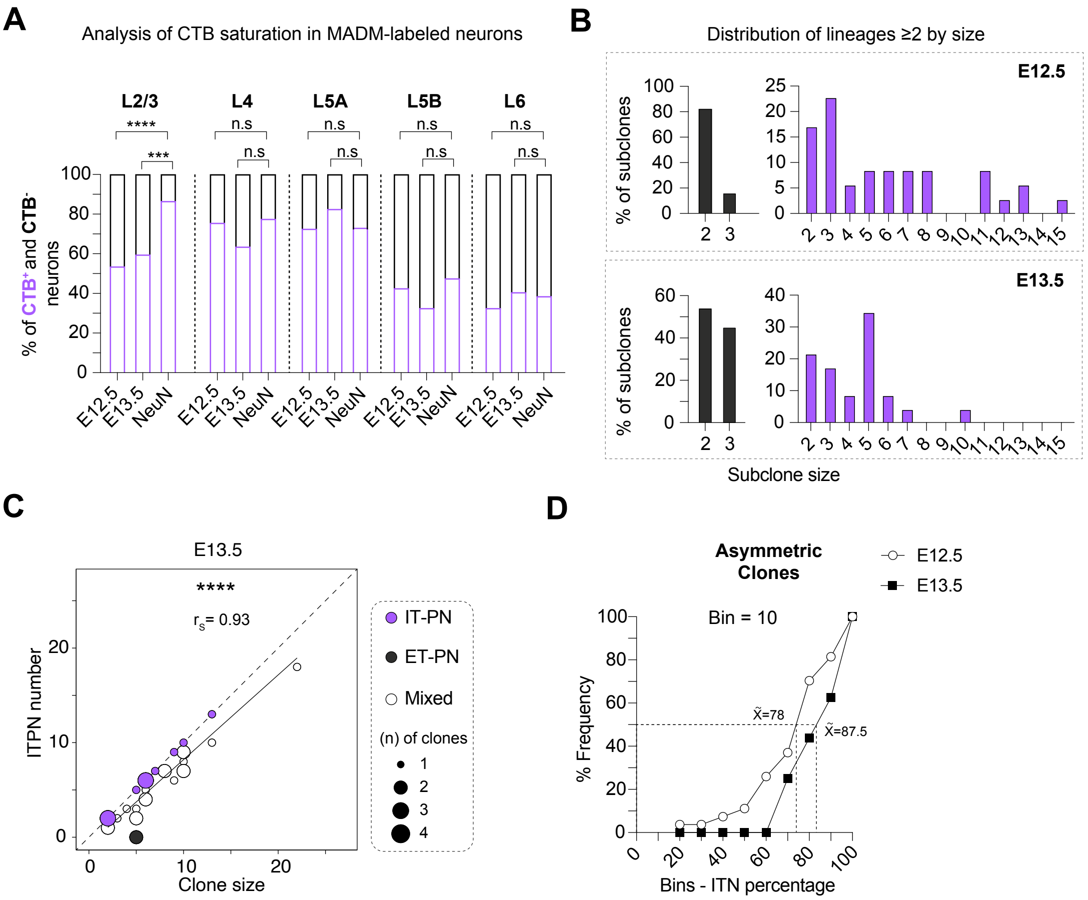


**Figure S6**_*Varela-Martínez et al.,*

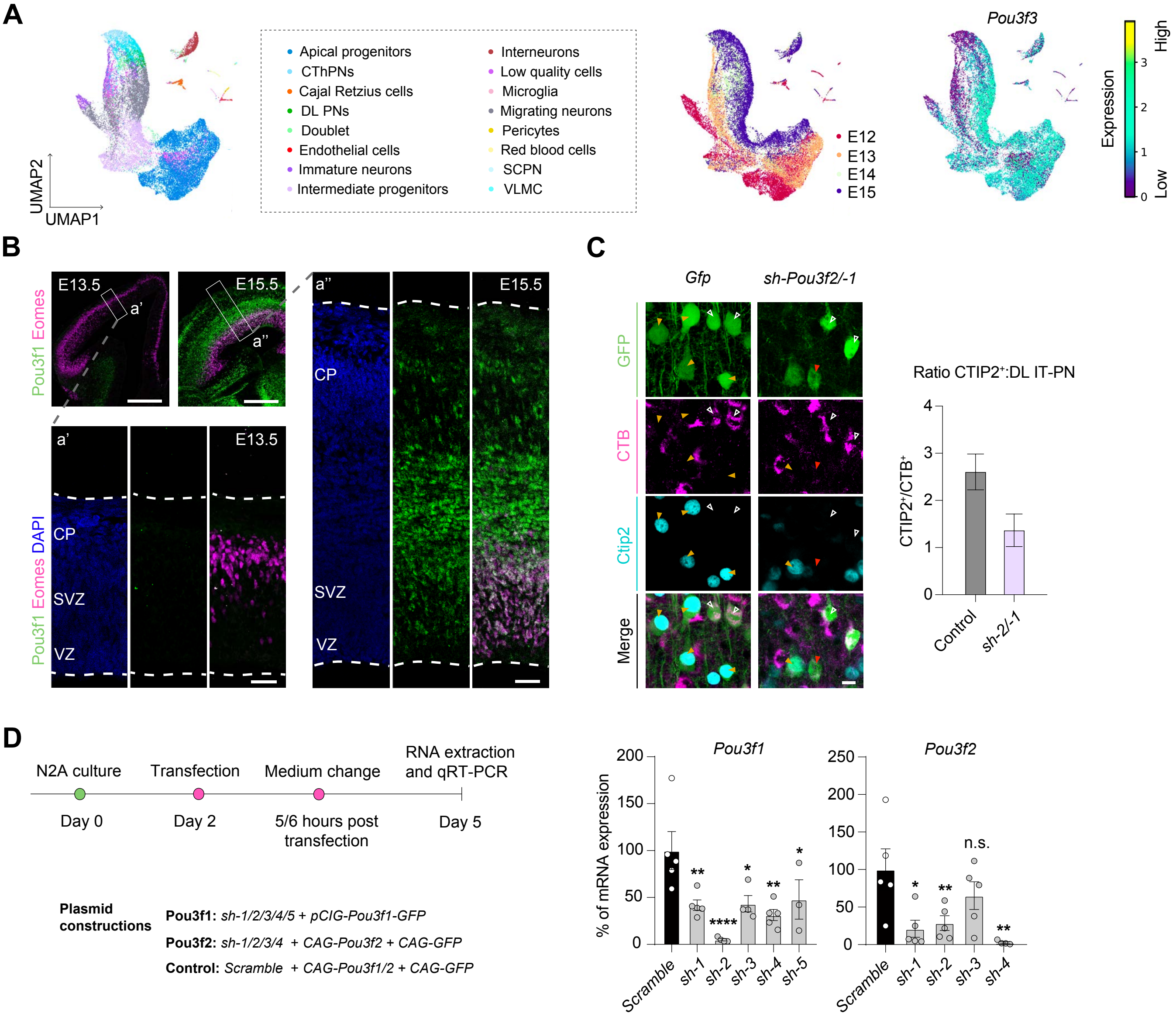


**Figure S7**_*Varela-Martínez et al.,*

### Atlas of MADM clones and subclones

Varela-Martínez et al.,

#### INDEX


**1.** Proliferative (P) E12.5

**2.** Proliferative (P) E13.5

**3.** Asymmetric (A) E12.5

**4.** Asymmetric (A) E13.5

**5.** Terminal (T) E12.5

**6.** Terminal (T) E13.5

**7.** Orphan (O) E12.3 and E13.5


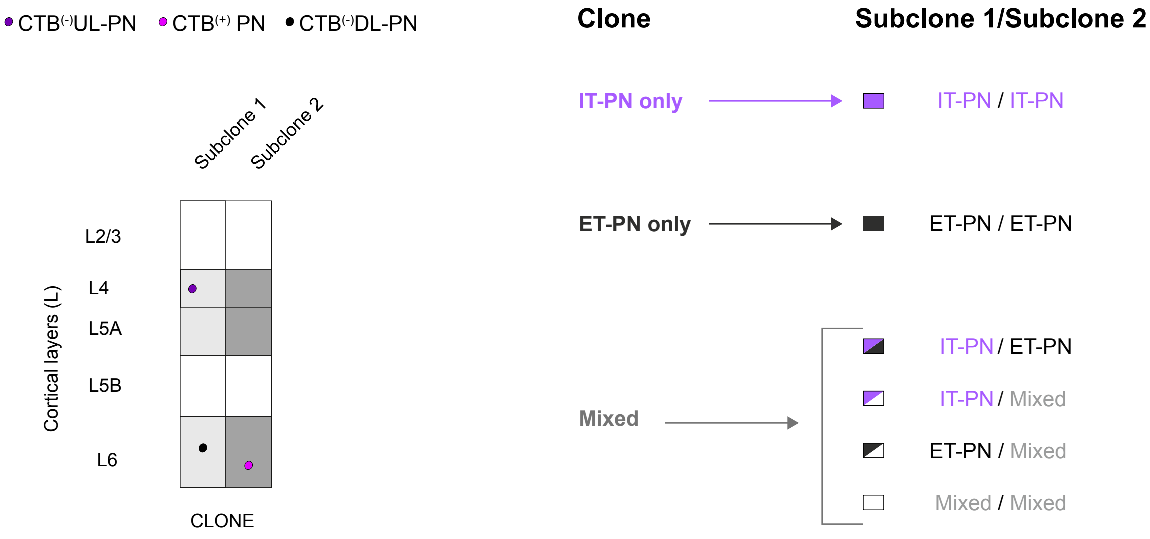

#### Proliferative clones E12.5: • CTB(-)UL-PN • CTB(+) PN • CTB(-)DL-PN

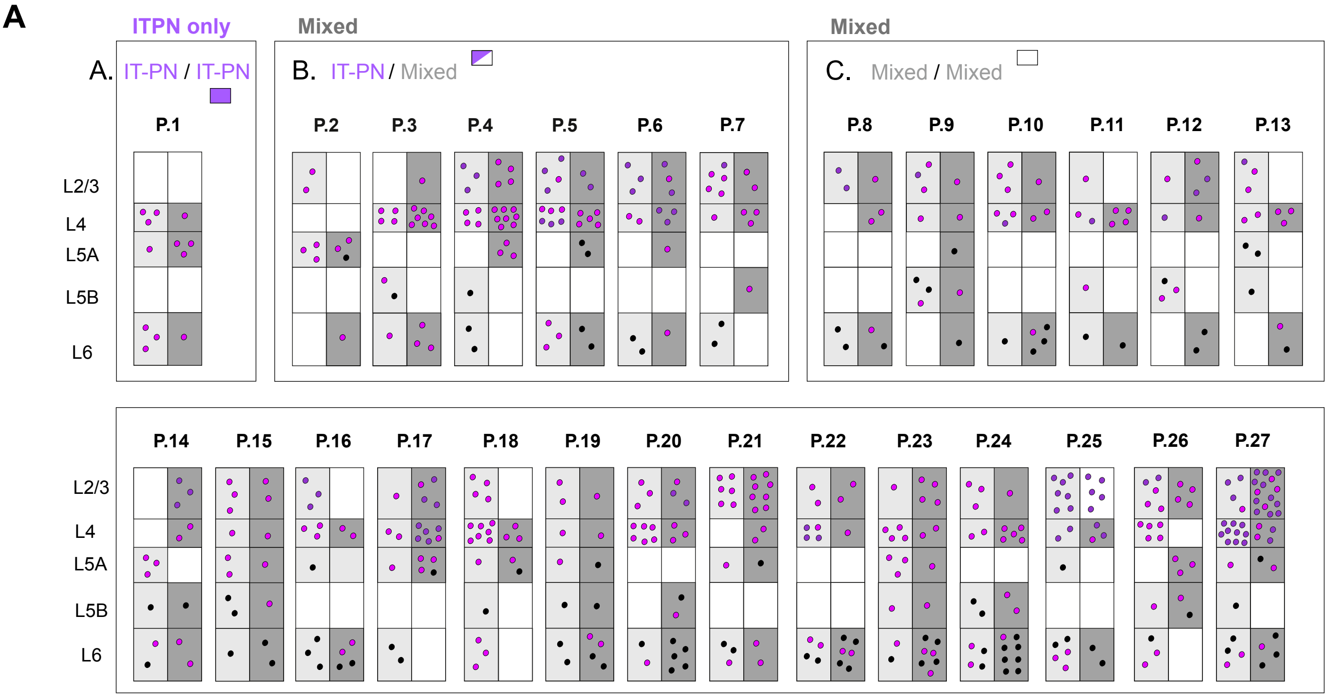


**B**

| E12.5 | | Proliferative clones Subclone.1 | Subclone.2 | Size .1 | Size .2 | Total Size |
|---|---|---|---|---|---|---|
| A. IT-PN/IT-PN | P.1 | L6 L5A L5A L5A L4 | L6 L6 L6 L5A L4 L4 L4 | 5 | 7 | 12 |
| B. IT-PN/Mixed | P.2 | L6 L5A L5A L5A | L5A L5A L5A L2/3 L2/3 | 4 | 5 | 9 |
| | P.3 | L5B L6 L5B L4 L4 L4 L4 | L6 L6 L6 L4 L4 L4 L4 L4 L4 L4 L2/3 | 7 | 11 | 18 |
| | P.4 | L6 L6 L5B L4 L4 L4 L4 L2/3 L2/3 L2/3 | L5A L5A L5A L4 L4 L4 L4 L4 L4 L4 L4 L2/3 L2/3 L2/3 L2/3 | 10 | 15 | 25 |
| | P.5 | L6 L6 L5A L5A L4 L4 L4 L4 L4 L2/3 L2/3 | L6 L6 L6 L4 L4 L4 L4 L4 L4 L2/3 L2/3 L2/3 L2/3 | 11 | 13 | 24 |
| | P.6 | L6 L6 L4 L4 L2/3 L2/3 L2/3 | L6 L5A L4 L4 L4 L2/3 L2/3 L2/3 | 7 | 8 | 15 |
| | P.7 | L6 L6 L4 L2/3 L2/3 L2/3 L2/3 L2/3 | L5B L4 L4 L4 L2/3 L2/3 | 8 | 6 | 14 |
| C. Mixed/Mixed | P.9 | L6 L6 L2/3 L2/3 | L6 L6 L4 L4 L2/3 | 4 | 5 | 9 |
| | P.8 | L6 L5A L5B L4 L2/3 | L5B L5B L5B L4 L2/3 L2/3 L2/3 | 5 | 7 | 12 |
| | P.10 | L6 L4 L4 L4 L2/3 L2/3 L2/3 | L6 L6 L6 L6 L4 L4 L2/3 | 7 | 7 | 14 |
| | P.11 | L6 L5B L4 L4 L2/3 | L6 L4 L4 L4 L4 | 5 | 5 | 10 |
| | P.12 | L5B L5B L5B L4 L2/3 | L6 L6 L4 L2/3 L2/3 L2/3 | 5 | 6 | 11 |
| | P.13 | L6 L6 L4 L4 L4 | L5B L5A L5A L4 L4 L2/3 L2/3 L2/3 | 5 | 8 | 13 |
| | P.14 | L6 L5B L6 L5A L5A L5A | L5B L6 L6 L4 L4 L2/3 L2/3 L2/3 | 6 | 8 | 14 |
| | P.15 | L6 L6 L5B L5A L4 L2/3 L2/3 | L6 L5B L5B L5A L5A L4 L2/3 L2/3 L2/3 | 7 | 9 | 16 |
| | P.16 | L6 L6 L6 L6 L4 L4 | L6 L6 L6 L5A L4 L4 L4 L2/3 L2/3 L2/3 | 6 | 10 | 16 |
| | P.17 | L6 L6 L5A L4 L4 L2/3 | L5A L5A L5A L5A L4 L4 L4 L4 L4 L4 L2/3 L2/3 L2/3 L2/3 | 6 | 14 | 20 |
| | P.18 | L5A L5A L4 L4 L4 | L5B L6 L6 L6 L5A L4 L4 L4 L4 L4 L4 L4 L2/3 L2/3 L2/3 L2/3 | 5 | 16 | 21 |
| | P.19 | L6 L6 L5B L5A L4 L2/3 L2/3 | L6 L6 L5B L6 L6 L5A L4 L2/3 | 7 | 8 | 21 |
| | P.20 | L6 L6 L4 L4 L4 L4 L4 L4 L2/2 L2/2 L2/3 | L6 L6 L6 L6 L6 L5B L5B L4 L4 L4 L2/3 L2/3 L2/3 | 11 | 13 | 24 |
| | P.21 | L6 L6 L6 L5A L2/3 L2/3 L2/3 L2/3 L2/3 L2/3 | L6 L6 L5A L4 L4 L2/3 L2/3 L2/3 L2/3 L2/3 L2/3 L2/3 L2/3 | 10 | 13 | 23 |
| | P.22 | L6 L6 L6 L6 L6 L6 L4 L2/3 L2/3 | L6 L6 L6 L6 L4 L4 L4 L4 L2/3 L2/3 | 9 | 10 | 19 |
| | P.23 | L6 L6 L5B L5A L5A L5A L4 L4 L4 L4 L2/3 | L6 L6 L6 L6 L6 L6 L6 L5B L5A L4 L4 L2/3 L2/3 L2/3 L2/3 | 11 | 15 | 26 |
| | P.24 | L6 L6 L5B L5B L6 L6 L4 L4 L2/3 L2/3 L2/3 | L6 L6 L6 L6 L6 L6 L6 L6 L5B L5B L4 L4 L4 L4 L4 L2/3 | 11 | 16 | 27 |
| | P.25 | L6 L6 L4 L4 L4 L2/3 L2/3 L2/3 L2/3 L2/3 | L6 L6 L6 L6 L6 L5A L4 L4 L2/3 L2/3 L2/3 L2/3 L2/3 L2/3 L2/3 | 10 | 15 | 25 |
| | P.26 | L5B L5B L5A L5A L5A L2/3 L2/3 L2/3 L2/3 | L6 L6 L6 L5B L4 L4 L4 L4 L4 L2/3 L2/3 L2/3 L2/3 L2/3 | 9 | 14 | 23 |
| | P.27 | L6 L6 L5B L6 L6 L5A L4 L4 L4 L4 L4 L4 L4 L4 L4 L4 L2/3 L2/3 L2/3 L2/3 | L6 L6 L6 L6 L5A L5A L4 L4 L4 L4 L2/3 L2/3 L2/3 L2/3 L2/3 L2/3 L2/3 L2/3 L2/3 L2/3 L2/3 L2/3 L2/3 L2/3 L2/3 | 20 | 25 | 45 |

#### Proliferative clones E13.5: • CTB$^{(-)}$UL-PN • CTB$^{(+)}$ PN • CTB$^{(-)}$DL-PN

**A**

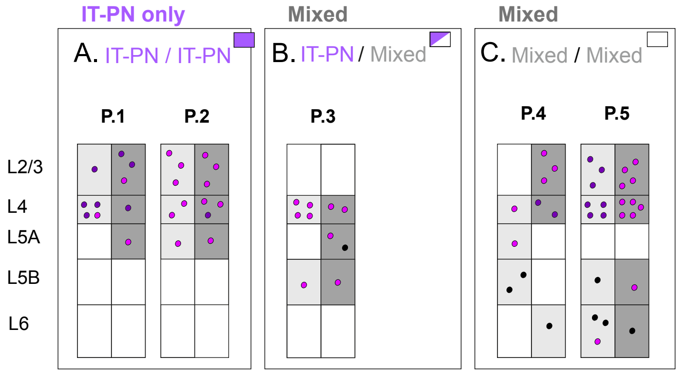


**B**

| E13.5 | | Proliferative clones Subclone 1 | Subclone 2 | Size .1 | .2 | Total Size |
|---|---|---|---|---|---|---|
| A. IT-PN/IT-PN | P.1 | L4 L4 L4 L4 L2/3 | L5A L4 L2/3 L2/3 L2/3 | 5 | 5 | 10 |
| | P.2 | L5A L4 L4 L2/3 L2/3 L2/3 | L5A L4 L4 L4 L2/3 L2/3 L2/3 | 6 | 7 | 13 |
| B. IT-PN/Mixed | P.4 | L5B L4 L4 L4 L4 | L5B L5A L5A L4 L4 | 4 | 6 | 10 |
| C. Mixed/Mixed | P.4 | L5B L5B L5A L4 | L6 L4 L4 L2/3 L2/3 L2/3 | 4 | 6 | 10 |
| | P.5 | L6 L6 L6 L5B L4 L4 L4 L4 L2/3 L2/3 L2/3 | L6 L5B L4 L4 L4 L4 L4 L2/3 L2/3 L2/3 L2/3 | 6 | 7 | 13 |

#### Asymmetric clones E12.5: • CTB(-)UL-PN • CTB(+) PN • CTB(-)DL-PN

**A**

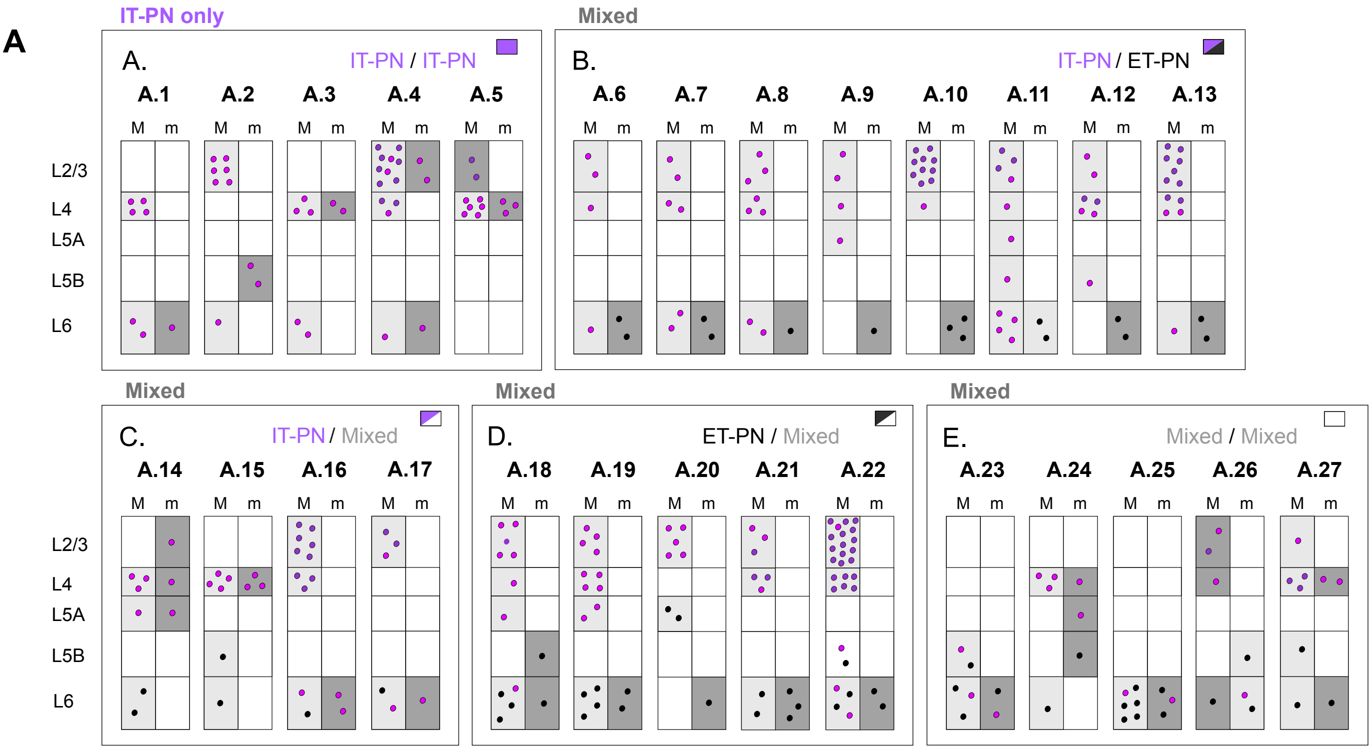


**B**

| E12.5 | | minority (m) | MAJORITY (M) Asymmetric clones | Size m | Size M | Size |
|---|---|---|---|---|---|---|
| A. IT-PN/IT-PN | A.1 | L6 | L6 L6 L4 L4 L4 L4 | 1 | 6 | 7 |
| | A.2 | L5B L5B | L6 L2/3 L2/3 L2/3 L2/3 L2/3 L2/3 | 2 | 7 | 9 |
| | A.3 | L4 L4 | L6 L6 L4 L4 L4 | 2 | 5 | 7 |
| | A.4 | L6 L2/3 L2/3 | L6 L4 L4 L4 L2/3 L2/3 L2/3 L2/3 L2/3 L2/3 L2/3 L2/3 L2/3 | 3 | 13 | 16 |
| | A.5 | L4 L4 L4 | L4 L4 L4 L4 L4 L4 L2/3 L2/3 | 3 | 8 | 11 |
| B. ITP-N/ET-PN | A.6 | L6 L6 | L6 L4 L2/3 L2/3 | 2 | 4 | 6 |
| | A.7 | L6 L6 | L6 L6 L4 L4 L2/3 L2/3 | 2 | 6 | 8 |
| | A.8 | L6 | L6 L6 L4 L4 L4 L2/3 L2/3 L2/3 | 1 | 8 | 9 |
| | A.9 | L6 | L5A L4 L2/3 L2/3 | 1 | 4 | 5 |
| | A.10 | L6 L6 L6 | L4 L2/3 L2/3 L2/3 L2/3 L2/3 L2/3 L2/3 L2/3 L2/3 L2/3 | 3 | 11 | 14 |
| | A.11 | L6 L6 | L6 L6 L6 L6 L5B L5A L4 L2/3 L2/3 L2/3 L2/3 | 2 | 11 | 13 |
| | A.12 | L6 L6 | L5B L4 L4 L4 L4 L2/3 L2/3 | 2 | 7 | 9 |
| | A.13 | L6 L6 | L6 L4 L4 L4 L4 L2/3 L2/3 L2/3 L2/3 L2/3 L2/3 L2/3 | 2 | 12 | 14 |
| C. IT-PN/Mixed | A.14 | L5A L4 L2/3 | L6 L6 L5A L4 L4 L4 | 3 | 6 | 9 |
| | A.15 | L4 L4 L4 | L6 L5B L4 L4 L4 L4 | 3 | 6 | 9 |
| | A.16 | L6 L6 | L6 L6 L4 L4 L4 L2/3 L2/3 L2/3 L2/3 L2/3 L2/3 | 2 | 11 | 13 |
| | A.17 | L6 | L6 L6 L2/3 L2/3 L2/3 | 1 | 5 | 6 |
| D. ET-PN/Mixed | A.18 | L6 L5B | L6 L6 L6 L6 L5A L4 L2/3 L2/3 L2/3 L2/3 L2/3 L2/3 | 2 | 11 | 13 |
| | A.19 | L6 L6 | L6 L6 L6 L6 L5A L5A L4 L4 L4 L4 L2/3 L2/3 L2/3 L2/3 | 2 | 14 | 16 |
| | A.20 | L6 | L5A L5A L2/3 L2/3 L2/3 L2/3 L2/3 | 1 | 7 | 8 |
| | A.21 | L6 L6 L6 | L6 L6 L4 L4 L4 L2/3 L2/3 L2/3 | 3 | 8 | 11 |
| | A.22 | L6 L6 | L6 L6 L5B L6 L6 L5B L4 L4 L4 L4 L4 L2/3 L2/3 L2/3 L2/3 L2/3 L2/3 L2/3 L2/3 L2/3 L2/3 L2/3 L2/3 L2/3 L2/3 L2/3 L2/3 | 2 | 27 | 29 |
| E. Mixed/Mixed | A.23 | L6 L6 | L6 L6 L5B L6 L5B | 2 | 5 | 7 |
| | A.24 | L5B L5A L4 | L6 L4 L4 L4 | 3 | 4 | 7 |
| | A.25 | L6 L6 L6 | L6 L6 L6 L6 L6 L6 | 3 | 6 | 9 |
| | A.26 | L6 L5B L6 | L6 L4 L2/3 L2/3 | 3 | 4 | 7 |
| | A.27 | L6 L4 L4 | L6 L5B L4 L4 L4 L2/3 | 3 | 6 | 9 |

#### Asymmetric clones E13.5: • CTB(-)UL-PN • CTB(+) PN • CTB(-)DL-PN

**A**

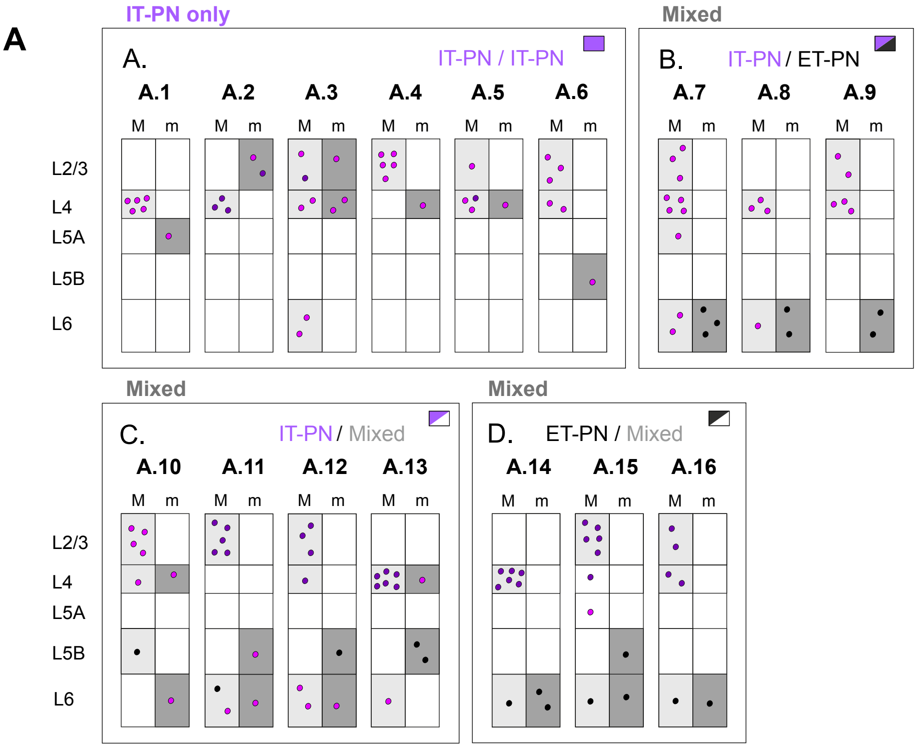


**B**

| E13.5 | | minority (m) | MAJORITY (M) — Asymmetric clones | Size m | Size M | Size |
|---|---|---|---|---|---|---|
| A. IT-PN/IT-PN | A.1 | L5A | L4 L4 L4 L4 L4 | 1 | 5 | 6 |
| | A.2 | L2/3 L2/3 | L4 L4 L4 L2/3 L2/3 | 2 | 5 | 7 |
| | A.3 | L4 L4 L2/3 | L6 L6 L4 L4 L2/3 L2/3 | 3 | 6 | 9 |
| | A.4 | L4 | L2/3 L2/3 L2/3 L2/3 L2/3 | 1 | 5 | 6 |
| | A.5 | L4 | L4 L4 L4 L2/3 | 1 | 4 | 5 |
| | A.6 | L5B | L4 L4 L2/3 L2/3 L2/3 | 1 | 5 | 6 |
| B. IT-PN/ET-PN | A.7 | L6 L6 L6 | L6 L6 L5A L4 L4 L4 L4 L2/3 L2/3 L2/3 | 3 | 10 | 13 |
| | A.8 | L6 L6 | L6 L4 L4 L4 | 2 | 4 | 6 |
| | A.9 | L6 | L4 L4 L4 L2/3 L2/3 | 1 | 5 | 6 |
| C. IT-PN/Mixed | A.10 | L4 L6 | L5B L4 L2/3 L2/3 L2/3 L2/3 | 2 | 6 | 8 |
| | A.11 | L6 L5B | L6 L6 L2/3 L2/3 L2/3 L2/3 L2/3 L2/3 | 2 | 8 | 10 |
| | A.12 | L5B L6 | L6 L6 L4 L2/3 L2/3 L2/3 | 2 | 6 | 8 |
| | A.13 | L5B L5B L4 | L6 L4 L4 L4 L4 L4 L4 | 3 | 7 | 10 |
| D. ET-PN/Mixed | A.14 | L6 L6 | L6 L4 L4 L4 L4 L4 L4 | 2 | 7 | 9 |
| | A.15 | L6 L5B | L6 L5A L4 L2/3 L2/3 L2/3 L2/3 L2/3 | 2 | 8 | 10 |
| | A.16 | L6 | L6 L4 L4 L2/3 L2/3 | 1 | 5 | 6 |

#### Terminal clones E12.5: • CTB(-)UL-PN • CTB(+) PN • CTB(-)DL-PN

A

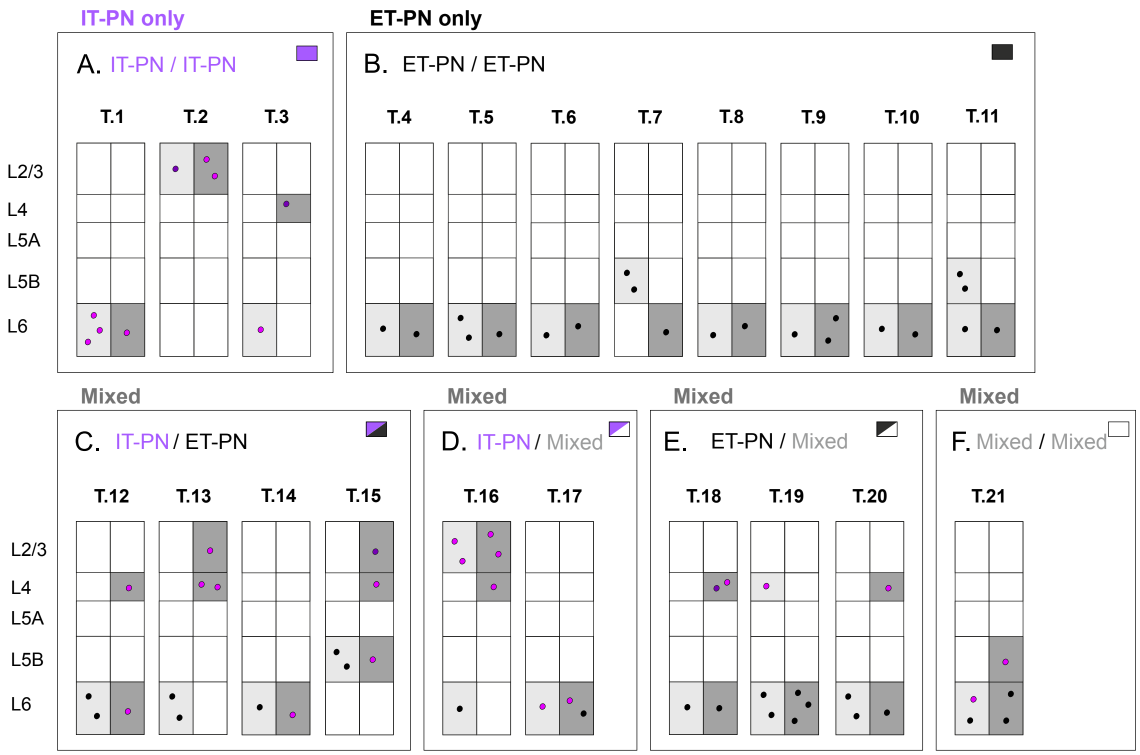


B

| E12.5 | | Subclone 1 | Subclone 2 | Size .1 | Size .2 | Size |
|---|---|---|---|---|---|---|
| A. IT-PN/IT-PN | T.1 | L6 L6 L6 | L6 | 3 | 1 | 4 |
| | T.2 | L2/3 | L2/3 L2/3 | 1 | 2 | 3 |
| | T.3 | L6 | L4 L4 | 1 | 2 | 3 |
| B. ET-PN/ET-PN | T.4 | L6 | L6 | 1 | 1 | 2 |
| | T.5 | L6 L6 | L6 | 2 | 1 | 3 |
| | T.6 | L5B L5B | L6 | 2 | 1 | 3 |
| | T.7 | L6 | L6 | 1 | 1 | 2 |
| | T.8 | L6 | L6 L6 | 1 | 2 | 3 |
| | T.9 | L6 | L6 | 1 | 1 | 2 |
| | T.10 | L6 | L6 | 1 | 1 | 2 |
| | T.11 | L6 L5B L5B | L6 | 3 | 1 | 4 |
| C. IT-PN/ET-PN | T.12 | L6 L6 | L6 L4 | 2 | 2 | 4 |
| | T.13 | L6 L6 | L4 L4 L2/3 | 2 | 3 | 5 |
| | T.14 | L6 | L6 | 1 | 1 | 2 |
| | T.15 | L5B L5B | L5B L4 L2/3 | 2 | 3 | 5 |
| D. IT-PN/Mixed | T.16 | L4 L2/3 L2/3 | L6 L2/3 L2/3 | 3 | 3 | 6 |
| | T.17 | L6 | L6 L6 | 1 | 2 | 3 |
| E. ET-PN/Mixed | T.18 | L6 | L6 L4 L4 | 1 | 3 | 4 |
| | T.19 | L6 L6 L6 | L6 L6 L4 | 3 | 3 | 6 |
| | T.20 | L6 L6 | L6 L4 | 2 | 2 | 4 |
| F. Mixed/Mixed | T.21 | L6 L6 | L6 L6 L5B | 2 | 3 | 5 |

Header note: "Terminal clones" spans the Subclone 1 and Subclone 2 columns.

**Terminal clones E13.5:** • CTB(-)UL-PN • CTB(+) PN • CTB(-)DL-PN

**A**

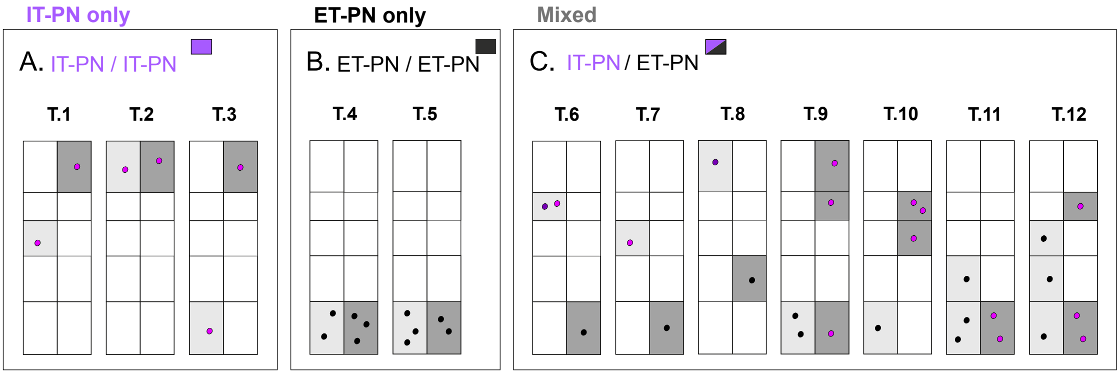


**B**

| E13.5 | | Subclone1 | Subclone 2 | Terminal clones | Size .1 | Size .2 | Size |
|---|---|---|---|---|---|---|---|
| A. IT-PN/IT-PN | T.1 | L5A | L2/3 | | 1 | 1 | 2 |
| | T.2 | L2/3 | L2/3 | | 1 | 1 | 2 |
| | T.3 | L6 | L2/3 | | 1 | 1 | 2 |
| B. ET-PN/ET-PN | T.4 | L6 L6 | L6 L6 L6 | | 2 | 3 | 5 |
| | T.5 | L6 L6 L6 | L6 L6 | | 3 | 2 | 5 |
| C. IT-PN/ET-PN | T.6 | L4 L4 | L6 | | 2 | 1 | 3 |
| | T.7 | L5A | L6 | | 1 | 1 | 2 |
| | T.8 | L2/3 | L5B | | 1 | 1 | 2 |
| | T.9 | L6 L4 L2/3 | L6 L6 | | 3 | 2 | 5 |
| | T.10 | L5A L4 L4 | L6 | | 3 | 1 | 4 |
| | T.11 | L6 L6 | L6 L6 L5B | | 2 | 3 | 5 |
| | T.12 | L6 L6 L4 | L6 L5B L5A | | 3 | 3 | 6 |

#### Orphan clones E12.5: • CTB(-)UL-PN • CTB(+) PN • CTB(-)DL-PN

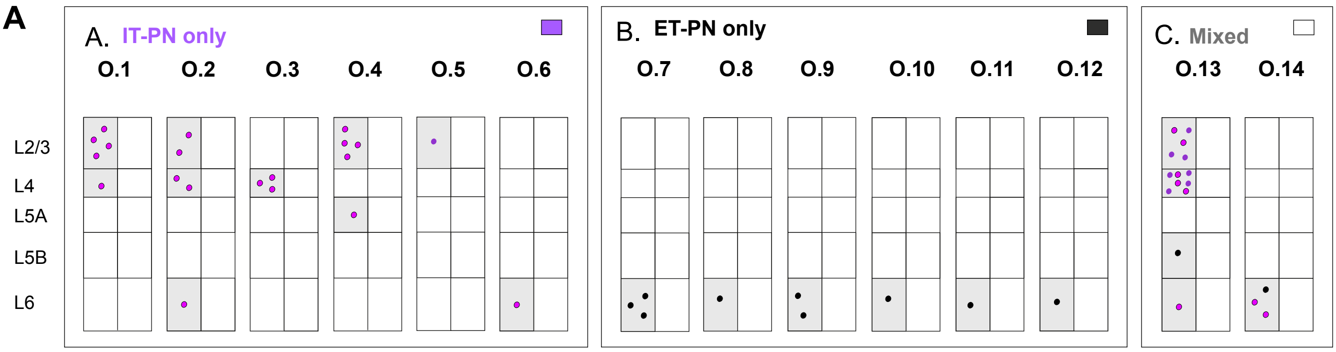


**B** **E12.5**

| | | Orphan clones | Size |
|---|---|---|---|
| A. (ITPN) | O.1 | L4 L2/3 L2/3 L2/3 L2/3 | 5 |
| | O.2 | L6 L4 L4 L2/3 L2/3 | 5 |
| | O.3 | L4 L4 L4 | 3 |
| | O.4 | L5A L2/3 L2/3 L2/3 L2/3 | 5 |
| | O.5 | L2/3 | 1 |
| | O.6 | L6 | 1 |
| B. (ETPN) | O.9 | L6 L6 L6 | 3 |
| | O.10 | L6 | 1 |
| | O.11 | L6 L6 | 2 |
| | O.12 | L6 | 1 |
| | O.13 | L6 | 1 |
| | O.14 | L6 | 1 |
| C. (Mixed) | O.7 | L6 L5B L4 L4 L4 L4 L4 L4 L4 L2/3 L2/3 L2/3 L2/3 L2/3 | 14 |
| | O.8 | L6 L6 L6 | 3 |

**C** **E13.5**

| | | Orphan clones | Size |
|---|---|---|---|
| A. (ITPN) | O.1 | L4 L2/3 L2/3 | 3 |
| | O.2 | L6 | 1 |
| | O.3 | L4 L2/3 | 2 |
| | O.4 | L4 L2/3 L2/3 L2/3 | 4 |
| | O.5 | L4 | 1 |
| | O.6 | L2/3 | 1 |
| | O.7 | L4 L2/3 | 2 |
| | O.8 | L2/3 L2/3 L2/3 L2/3 L2/3 L2/3 L2/3 L2/3 | 8 |
| | O.9 | L4 L4 | 2 |
| | O.10 | L2/3 | 1 |
| | O.11 | L4 L4 L4 L2/3 L2/3 | 5 |
| | O.12 | L5A L4 L2/3 L2/3 L2/3 | 5 |
| | O.13 | L2/3 | 1 |
| | O.14 | L6 L5A L4 | 3 |
| B. (ETPN) | O.15 | L5A | 1 |
| | O.16 | L5A | 1 |

#### Orphan clones E13 .5: • CTB(-)UL-PN • CTB(+) PN • CTB(-)DL-PN

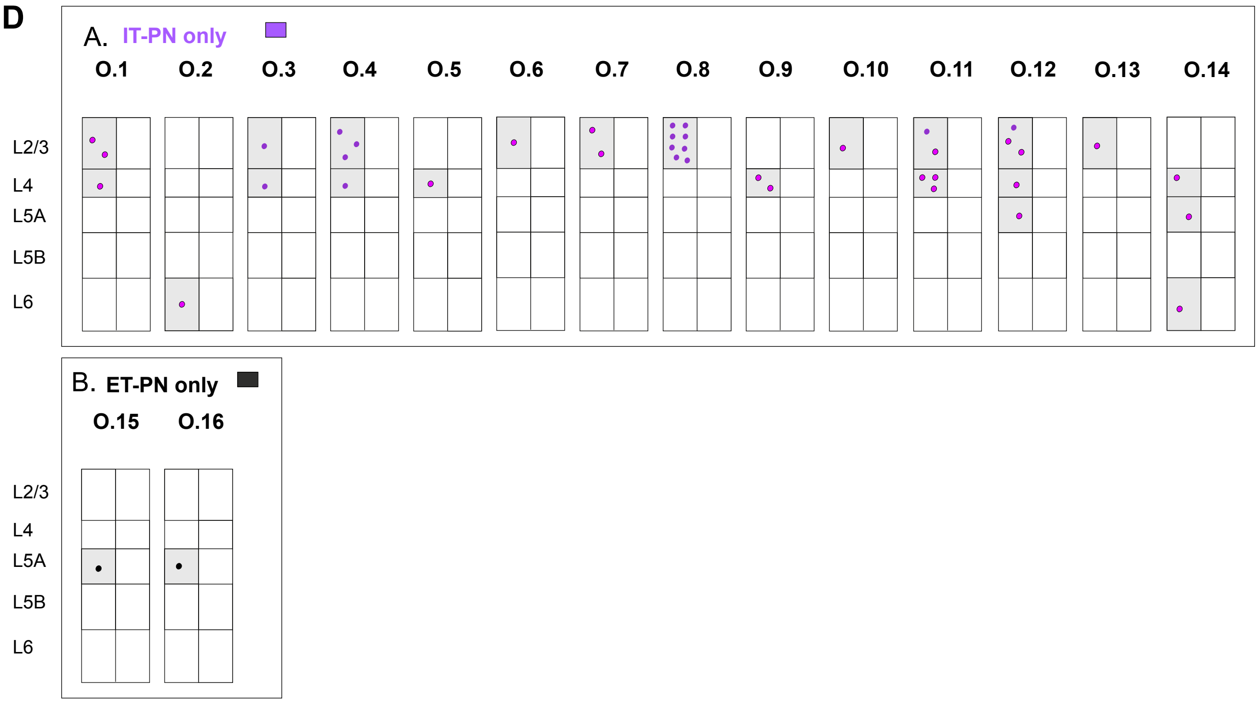
